# NADPH Oxidase 2 Derived Reactive Oxygen Species Promote CD8^+^ T cell Effector Function

**DOI:** 10.1101/2021.02.24.432756

**Authors:** Jing Chen, Chao Liu, Anna V. Chernatynskaya, Brittney Newby, Todd M. Brusko, Yuan Xu, Nadine Morgan, Christopher P. Santarlas, Westley H. Reeves, Hubert M. Tse, Jennifer W. Leiding, Clayton E. Mathews

**Affiliations:** Department of Pathology, Immunology and Laboratory Medicine Gainesville, FL 32610, USA; Department of Medicine, University of Florida, Gainesville, FL 32610, USA; Department of Microbiology, University of Alabama at Birmingham, Birmingham, AL 35294, USA; Department of Pediatrics, Division of Allergy and Immunology, University of South Florida; Johns Hopkins All Children’s Hospital, St. Petersburg, FL 33701, USA

## Abstract

Oxidants participate in lymphocyte activation and function. We previously demonstrated that eliminating the activity of NADPH oxidase 2 (NOX2) significantly impaired the effectiveness of autoreactive CD8^+^ cytotoxic T lymphocytes (CTL). However, the molecular mechanisms impacting CTL function remain unknown. Here, we studied the role of NOX2 in both non-obese diabetic (NOD) mouse and human CTL function. Genetic ablation or chemical inhibition of NOX2 in CTL significantly suppressed activation-induced expression of the transcription factor T-bet, the master transcription factor of the Tc1 cell lineage, and T-bet target effectors genes such as IFNγ and granzyme B. Inhibition of NOX2 in both human and mouse CTL prevented target cell lysis. We identified that superoxide generated by NOX2 must be converted into hydrogen peroxide to transduce the redox signal in CTL. Further, we show that NOX2-generated oxidants deactivate the Tumor Suppressor Complex leading to mTOR complex 1 activation and CTL effector function. These results indicate that NOX2 plays a non-redundant role in T cell receptor-mediated CTL effector function.

## INTRODUCTION

Cytotoxic CD8^+^ T lymphocytes (CTL) play vital roles in anti-tumor immunity, elimination of intracellular infections, and the pathogenesis of autoimmune diseases, including type 1 diabetes (T1D) (1, 2). Activation of CTL leads to cell expansion, differentiation, and downstream production of pro-inflammatory cytokines and cytotoxic factors, which are critical to induce cell death in antigen-bearing target cells. During CTL activation, multiple factors function collaboratively to control the master transcription factors that regulate effector function (3–5). Reactive oxygen species (ROS), a group of low molecular weight highly reactive molecules containing oxygen (6), have been proposed as one of the signal transducers for CTL activation and differentiation (7, 8).

At high concentrations, ROS can be toxic and participate in pathologies; however, regulated ROS production controls cellular functions by mediating redox signaling (6, 9, 10). Specific enzyme families, such as the NADPH oxidases (9), can modulate redox signal transduction through production of ROS. NOX2 is expressed in a wide variety of tissues/cell types and is the free radical source of the “respiratory burst” used by neutrophils and macrophages to destroy microbes and as a biological second messenger for signal transduction within lymphocytes (9, 10). We have reported that NOX2 regulates CD4^+^ T cell differentiation by promoting T_H_1 responses (11), a finding confirmed by subsequent studies with human samples (12). In addition, we have observed a CD8^+^ T cell intrinsic role of NOX2 that is essential for autoimmune diabetes initiation (8). Ergo, our data as well as those from other groups support an important role of NOX2 in adaptive immune function in addition to the well-established role of NOX2 in innate immunity (13, 14). The goal of the current study is to elucidate the mechanisms of NOX2 in regulating CD8^+^ T cell effector function.

T1D is caused by an autoimmune-mediated destruction of insulin-producing β cells (15–20). CTL are considered the final effector cells of β cell death in the non-obese diabetic (NOD) mouse (19, 21–24). CTL induce β cell death by utilizing pro-inflammatory cytokines and cytotoxic effector molecules such as IFNγ and TNFα, perforin, and granzyme B. We reported that NOX2 produced ROS contribute to the onset of diabetes in NOD mice using a NOD strain that encodes a non-functional NOX2 enzyme [NOD-*Ncf1^m1J^*] (8, 11, 25). Adoptive transfer of CTLs from NOX2-deficient NOD-*Ncf1^m1J^* donors into NOX2 intact recipients (NOD-*Scid*) resulted in a delayed onset of diabetes, indicating that NOX2 activity within the CTL is required for pathogenesis (8).

Production of pro-inflammatory cytokines and effector molecules as well as cytotoxic activity of human and mouse CTL are shown here to be blocked by genetic ablation or pharmacological inhibition of NOX2. Hydrogen peroxide derived from NOX2 promoted CTL effector function by enhancing the production of T-bet, the master transcription factor of the Tc1 cell lineage and driver of effectors genes such as IFNγ and granzyme B. ROS affected T-bet by regulating mTOR complex 1 (mTORc1) activity via Rheb-GTP levels through the redox sensitive Tsc1/Tsc2 signaling complex. Collectively, these findings define a mechanism by which NOX2 functions in CTL and highlights the significance of redox signaling in CTL-mediated diseases.

## MATERIALS AND METHODS

### Animals

NOD-*Ncf1^m1J^* mice, described previously (8, 11, 25), were bred and maintained at the University of Florida. All other mouse strains (NOD/ShiLtJ (NOD), NOD.CB17-*Prkdc^scid^*/J (NOD-*Scid*), NOD.Cg-*Rag1^tm1Mom^-*Tg (TcraAI4)^1Dvs^/DvsJ (NOD.AI4α-*Rag1^-/-^*), NOD.Cg-*Rag1^tm1Mom^*Tg (TcrbAI4)^1Dvs^/DvsJ (NOD.AI4β-*Rag1*^-/-^) as well as C57BL6/J (B6) mice) were purchased from The Jackson Laboratory (Bar Harbor, ME. F1 hybrid progeny were developed from outcrosses of NOD.AI4α-*Rag1^-/-^* to NOD.AI4β-*Rag1^-/-^* and referred to as NOD.AI4α/β-*Rag1^-/-^*. Female mice were used for all experiments. All mice used in this study were housed in specific pathogen free facilities and all procedures were approved by the University of Florida institution animal care and use committee.

### Human Subjects

The Institutional Review Board of the University of Florida approved all studies with human samples. Leukopaks from healthy donors (50% female, age 16-25 years, median 19) were purchased from Life South Blood Centers. PBMCs were isolated using Ficoll gradient method.

### Materials

Chemicals were purchased from Sigma-Aldrich. HRP conjugated secondary antibodies were purchased from Santa Cruz Biotechnology. Details of Fluorescently labeled antibodies used in flow cytometry analysis with Fortessa or Cytek Aurora are listed in Supplemental Material.

### Purification and Cell Culture of Mouse CTL

Spleens from age-matched NOD, NOD-*Ncf1^m1J^* and B6 females were collected, homogenized to a single cell suspension and subjected to hemolysis with Gey’s solution(26). CTL were negatively selected using paramagnetic beads following column selection (CD8^+^ T cell isolation kit (Miltenyi Biotec)). Purity was greater than 90% and confirmed on a BD LSR Fortessa (BD Biosciences). Purified CTL were cultured in complete DMEM. All cultures using mouse T cells were performed in complete DMEM (Low glucose (5.5mM) DMEM (Corning Cellgro) containing 10% fetal bovine serum (FBS, HyClone Laboratories), HEPES buffer, gentamicin (Gemini), MEM Non-Essential Amino Acids Solution (Life Technologies) and 2-mercaptoethanol.

### Immuno-Spin Trapping and Immunofluorescence

Macromolecular-centered free radicals were detected after stimulation of purified NOD and NOD.*Ncf1^m1J^* CD8^+^ T cells with α-CD3 (clone 145-2C11 (BD Pharmingen); 1μg/mL) and α-CD28 (clone 37.51(BD Pharmingen); 1μg/mL) in the presence of DMPO (1 mmol/L) in tissue culture–treated chamber slides. After 24 hours, cells were fixed in 4% paraformaldehyde in PBS, permeabilized with 0.1% Triton X-100 + 0.1% Tween in PBS, blocked with 10% BSA in PBS, and incubated with chicken IgY anti-DMPO (20μg/mL) and rat IgY anti-mouse CD8a (1:250, SouthernBiotech). DMPO adducts were detected with Alexa Fluor 488–conjugated goat anti-chicken IgY secondary antibody (1:400; Jackson ImmunoResearch). CD8^+^ T cells were identified with Cy3-conjugated donkey anti-rat secondary antibody (1:500; Jackson ImmunoResearch). Images were obtained with an Olympus IX81 Inverted Microscope using a 403 objective and analyzed with cellSens Dimension imaging software, version 1.9. For quantitation of fluorescence intensity, three to six images were obtained for each data point. Each image was collected at the same exposure time, adjusted to the same intensity level for standardization, and the fluorescence intensity was measured using ImageJ software (National Institutes of Health).

### CTL proliferation

CD8^+^ T cells (1 × 10^5^) were used for proliferation after stimulation with plate-bound α-CD3ε (0.1 μg/mL) and α-CD28 (1 μg/mL) or beads conjugated with α-CD3ε and α-CD28 in the presence or absence of 200µM apocynin (ED Millipore). This dose of apocynin was used based on dose response curves using increasing concentrations of apocynin (0-300µM) and analyzing viability by flow cytometry (Live-Dead Near IR (ThermoFisher), Supplemental Figure 2A) and IFNγ levels by ELISA (BD Biosciences) (Supplemental Figure 2B). Cell proliferation was measured by [^3^H]-thymidine (TdR) incorporation as described previously (11).

### Detection of cytokine production and secretion by mouse CTL

Purified CD8^+^ T cells (5 × 10^5^) from NOD, NOD-*Ncf1^m1J^*, and B6 mice were activated with plate bounded α-CD3ε and α-CD28 or phorbol 12-myristate 13-acetate (PMA; 50ng/mL) plus ionomycin (1µg/mL) for a total of 72 hours, as previously described (27, 28). Cells were stained with BV421-labeled α-CD8 and fixed then permeabilized prior to being stained with fluorescently labeled antibodies to IFNγ, Granzyme B, and T-bet. Data were collected using a BD LSR Fortessa by gating on CD8^+^ T cells and then producing histograms of the intracellular targets. Isotype controls and fluorescence minus one (FMO) conditions were used to set gates for marker positivity for the protein of interest. Results were analyzed by Flowjo 7.6.1-10.5.0 software.

For antigen-specific activation of mouse AI4 T cells, NOD.AI4α/β-*Rag1*^-/-^ splenocytes were activated with 0.1 µM mimotope (amino acid sequence YFIENYLEL) and 25U/mL IL-2 for 72 hours (29, 30) with or without the presence of 200 µm apocynin. Cells were stained with LIVE/DEAD™ Fixable Near-IR to exclude dead cells, blocked with Fc-block (BD Biosciences), and stained for surface markers CD3-PE-Cy5 (145-2C11, eBioscience), CD8-BV650 (53-6.7, Biolegend), CD62L-BV711 (MEL-14, Biolegend) and CD69-PerCP-Cy5.5 (H1.2F3, BD Biosciences). Then cells were fixed and permeabilized using eBioscience™ FoxP3 / Transcription Factor Fixation/Permeabilization kit, and stained for p-S6K-PE (clone #215247, R&D Systems), Perforin-APC (S16009A, Biolegend), IFNγ-FITC (XMG1.2, BD Biosciences), Granzyme B-PacBlue (GB11, Biolegend) and T-bet-BV605 (4B10, Biolegend). Data were analyzed on Aurora. Results were expressed as median fluorescence intensity (MFI) ratio to activated when gated on live CD3^+^CD8^+^.

Secreted IFNγ was analyzed using CD8^+^ T cells (5 × 10^4^) that were stimulated with α-CD3ε and α-CD28 conjugated beads (Life Technologies). IFNγ was detected with the BD OptEIA™ Mouse IFNγ ELISA Set (BD Biosciences). ELISA plates were read on a SpectraMax Multi-Mode Microplate Reader and analyzed using Softmax Pro v.5.0.1 (Molecular Devices Corp.).

### Quantitative Real-Time PCR (qRT-PCR)

Total RNA from purified CD8^+^ T cells was isolated with TRIzol reagent (Invitrogen). cDNA was prepared using the Superscript III First-Strand Synthesis System (Invitrogen) according to the manufacturer’s protocol. Real-time quantitative PCR was performed as previously reported (31–33) using SYBR Green I (Bio-Rad) on a Roche LightCycler 480 II. The amplification program utilized the following steps for all primer sets: 95°C for 10 min, followed by 45 cycles of 95°C for 30s, 60°C for 30s and 72°C for 30s. The melting-curves of the PCR products were measured to ensure the specificity. Primers (Supplemental Table) were designed using qPrimerDepot (NIH).

### Mouse Cell-Mediated Lymphocytoxicity (CML) Assays

CML assays were performed as described previously (16, 18, 34, 35). Briefly, splenocytes from NOD.AI4α/β-*Rag1*^-/-^ mice were primed in RPMI1640 complete media containing 25 U/mL IL-2 and 0.1μM mimotope (amino acid sequence YFIENYLEL) for 3 days with (CTL Activation phase (+)) or without (CTL Activation phase (−)) 200μM apocynin. NOD-derived NIT-1 β cells (36) were plated at 10,000 cells per well in flat-bottom 96 well plates. NIT-1 cells were primed with 1000U/mL IFNγ for 24 hours. The NIT-1 cells were then labeled with 1 μCi/well of ^51^Cr (Perkin Elmer) for 3 hours at 37°C and co-cultured with AI4 T cells at a E:T ratio of 20:1 for 16 hours, with (CTL Effector phase (+)) or without (CTL Effector phase (−)) the presence of 200 μmM apocynin. Specific lysis was calculated as described (16, 18, 34, 35).

### Detection of cytokine production by human CD8^+^ T cells

Purified total human T cells were cultured in complete RPMI. Culture media contained the indicated concentration of apocynin. Total T cells (1 × 10^6^) were activated with plate bound α-CD3ε (OKT3, Biolgend) plus α-CD28 (CD28.2 RUO, BD Biosciences) at 5µg/mL each or corresponding isotype controls for a total of 72 hours. For IFNγdetection by ELISA, media supernatants were collected at 24, 48 and 72 hours and the IFNγ levels determined using BD OptEIA Human IFNγ ELISA set. For intracellular staining, cells were activated for 66 hours and then, re-stimulated with PMA (50ng/mL) plus ionomycin (1µg/mL) with the addition of GolgiStop (BD Biosciences) for 6 hours at 37°C in a 5% CO_2_ humid air incubator. Cell proliferation was measured by [^3^H]TdR incorporation. In independent cultures, cells were stained with α-human CD8-FITC (G42-8, BD Pharmingen), CD69-BV605 (FN50; Biolegend), CD4-PE-Cy7 (RPA-T4; eBiosciences) followed by intracellular staining to detect IFNγ - PerCP-Cy5.5 (4S.B3; Biolegend), granzyme B-AF647 (GB11; Biolegend), T-bet-Pacific Blue (4B10, Biolegend) and EOMES-PE (WD1928; eBiosciences). Intracellular staining was performed with the True Nuclear Transcription Factor staining kit (Biolegend) according to manufacturer’s protocol. Fluorescence was measured using BD LSR Fortessa and results analyzed using Flowjo 7.6.1 software.

Leukopak PBMC were pretreated with Apocynin 400uM, DMSO 0.04%, or left untreated in cRPMI for 1 hr at 37C in a CO2 incubator. Cells were then activated with soluble α-CD3 (1ug/mL) and α-CD28 (5ug/mL) or isotype antibodies, together with protein G (Sigma; 5ug/mL) as crosslinker for 10 min at 37°C in a CO2 incubator. At the end of activation, the reaction was stopped by immediately adding 4X Fix stock, to reach 1X concentration of Fix buffer and fixed for 30 min. Samples were then stained for viability with LIVE/DEAD™ Fixable Yellow Dead Cell Stain Kit (Invitrogen), blocked with Fc-block (ThermoFisher), permeabilized with ice-cold 90% methanol, and stained for surface markers CD3-eFluor450 (UCHT1, eBioscience), CD4-FITC (RPA-T4, Biolegend), CD8-BV711 (SK1, Biolegend) as well as intracellular molecules Phospho-S6 (Ser235, Ser236)-PE (cupk43k, ThermoFisher) and Phospho-AKT1 (Ser473)-APC (SDRNR, ThermoFisher). Data were analyzed on Cytek Aurora and results expressed as MFI ratio to activated when gated on Live/Dead_Yellow^-^ CD3^+^CD4^-^CD8^+^.

### CML Assays Using Auto-reactive Human CD8+ T cell Avatars and Human Derived BetaLox5 Cells as Target Cells

CD8^+^ T cells were pre-enriched from LeukoPaks by negative selection (RosetteSep (StemCell Technologies)) and then stained with CD8-APC/Cy7 (Clone SK1, BioLegend), CD45RA-PerCP/Cy5 (Clone H1100, BioLegend), and CD45RO-PE/Cy7 (Clone UCHL1, BioLegend). Naïve CD8^+^CD45RA^+^CD45RO^-^ T cells were sorted using a BD FACS Aria III. Purified (91.0 ± 2.5% purity) naïve CD8^+^ T cells were activated with α-CD3/CD28 Dynabeads (Life Technologies) in the presence or absence of Apocynin (200µM). At 48 hr, cells were transduced with a lentiviral vector containing an IGRP-specific, HLA-A*0201 restricted T Cell Receptor (TCR), as previously described (18). After transduction, CD8^+^ avatars were supplemented with complete RPMI (Corning Cellgro) and IL-2 (300U/mL) with or without apocynin (200µM) every 48 hours. Transduction efficiency for the cells expanded in apocynin was 32.4% whereas those expanded in media free of apocynin were transduced at 22.3%. After transduction cells were expanded for an additional 7 days and on day 9, the cells were cryopreserved in freeze medium (90% FBS/10% DMSO). Cells were thawed prior to and immediately used for the CML assays. CML assays were performed as described previously using BetaLox5 (BL5) cells as target (18, 34).

### Measurement of mTOR Complex Activity

mTOR1 complex activity analysis was preformed using purified CD8^+^ T cells (1 × 10^7^) from NOD and NOD-*Ncf1^m1J^* mice. Western blotting was used as the output assay to determine levels of the ribosomal protein S6 kinase (p-S6K) using an S6K and p-S6K antibodies (Cell Signaling Technology). To assess the impact of mTOR1 inhibition rapamycin (20ng/mL) treatment was performed 15 min prior to polyclonal activation of T cells. For redox reagent treatment, purified CD8^+^ T cells were exposed to either 5 μM phenylarsine oxide (PAO), 1 mM 2,3-dimercapto-1-propanol (British anti-Lewisite (BAL)), 10 μM H_2_O_2_, or 200 μM Apocynin 15 min before activation. All treatments were preformed in complete DMEM media. T cells were then activated with α-CD3ε and α-CD28 conjugated beads for 30 minutes. Cell lysates were then generated by sonication in RIPA buffer and subjected to western blot for p-S6K and total S6K. Lysates were separated on 7.5% (Bio-Rad) and transferred to 0.45-μm–charged PVDF membranes. Membranes were incubated overnight at 4°C with antibodies against phospho-S6K or total S6K and then exposed to the secondary antibody conjugated to HRP. For normalization, membranes were stripped and re-probed with α-Erk1, followed by incubation with HRP conjugated secondary antibody. Chemiluminescence was detected and photon signals converted to densitometry data with a FluorChem HD2 with AlphaView software (Alpha Innotech). Antigen-specific activation-induced p-S6K activity was also confirmed via flow cytometry, as noted above.

### Measurement of Akt activity and Rheb-GTP levels after T Cell Activation

Purified CD8^+^ T cells (2 × 10^7^) from NOD and NOD-*Ncf1^m1J^* mice were left unstimulated or subjected to stimulation with α-CD3ε and α-CD28 conjugated beads for 30 min. Akt kinase activity was measured using an Akt Kinase Activity Assay Kit (Cell Signaling Technology). Rheb-GTP levels were assessed using a Rheb Activation Assay Kit (New East Biosciences). p-AKT and p-S6 activity was also measured in human PBMC via flow cytometry.

### Statistics

Each experiment was run in triplicate with at least three independent trials. Statistical analysis was performed using GraphPad Prism (GraphPad Software) or SAS 9.2 (SAS Institute). Statistical significance between mean values was determined using the Student’s *t* test or one-way ANOVA with *P* < 0.05 considered as significantly different.

## RESULTS

### CD8^+^ T cells in NOD-*Ncf1^m1J^* Mice Exhibit a Reduced Production of Pro-inflammatory Cytokines and Effector Molecules

NOX2 functions in TCR signaling upon CD3 and CD28 cross-linking (8) (37), and ROS is produced during CD8^+^ T cell activation (38). ROS production by CD8^+^ T cells from NOD and NOD.*Ncf1^m1J^* mice was examined using the immuno-spin trap DMPO after stimulation with α-CD3 and α-CD28. Similar to our previous report (8), NOD.*Ncf1^m1J^* CD8^+^ T cells exhibited significantly less ROS production after stimulation when compared to CD8^+^ T cells from NOD mice (Supplemental Figure 1).

Earlier studies led us to hypothesize that NOX2-deficient NOD-*Ncf1^m1J^* CD8^+^ T cells would exhibit defective effector function *in vitro* (8). Stimulation of NOD-*Ncf1^m1J^* CD8^+^ T cells with α-CD3 and α-CD28 elicited a significant reduction in IFNγ production, which was consistent with our previous report that NOD-*Ncf1^m1J^* CD4^+^ T cells have decreased T_H_1 responses (11). TNFα and granzyme B were also reduced in the NOD-*Ncf1^m1J^* CD8^+^ T cells after α-CD3 and α-CD28 stimulation (Figure 1A). In addition, quantitative real-time PCR (qRT-PCR) was carried out with α-CD3/α-CD28 stimulated CTL. IFNγ and granzyme B transcripts were significantly reduced in NOD-*Ncf1^m1J^* CD8^+^ T cells (Figure 1B). No significant differences between NOD and NOD-*Ncf1^m1J^* CD8^+^ T cells were observed in the mRNA level of IL-2, IL-4, perforin, LAMP-1, TGFβ or FasL (Figure 1B). This indicated that a functional NOX2 is essential for the production of specific pro-inflammatory cytokines (IFNγ) and specific cytotoxic molecules (granzyme B). In accord with similar IL-2 transcription (Figure 1B), CD8^+^ T cell proliferation was equal in cells with and without NOX2 activity (Figure 1C). Differences in activation status, measured as CD8^+^CD62L^-^CD69^+^ cells (not shown), or apoptosis (Figure 1D) were not observed when comparing α-CD3 and α-CD28 stimulated NOD and NOD-*Ncf1^m1J^* CD8^+^ T cells. These data indicate the loss of ROS production in CTL resulted in a selective deficiency in a Tc1 effector phenotype.

**Figure 1.**
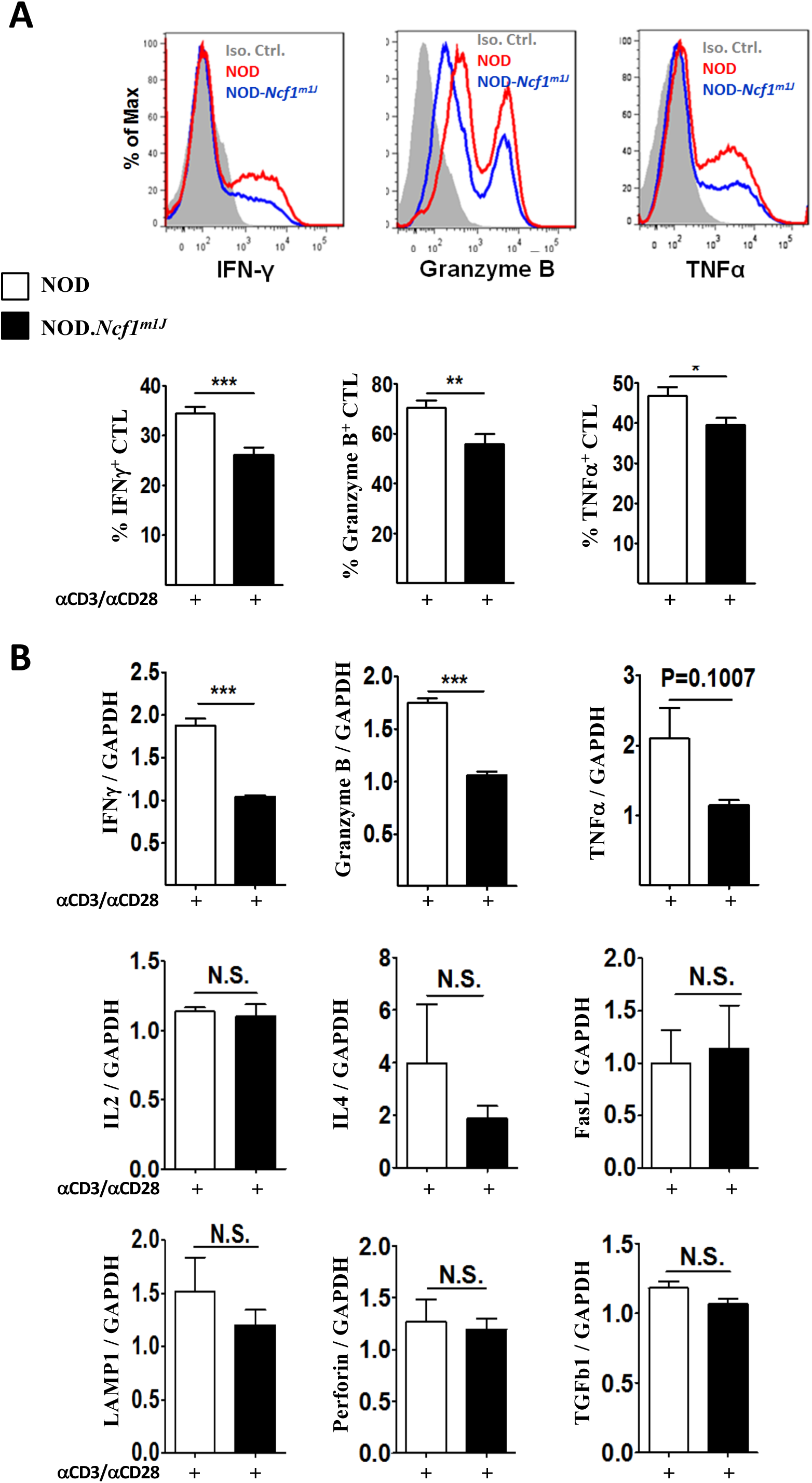
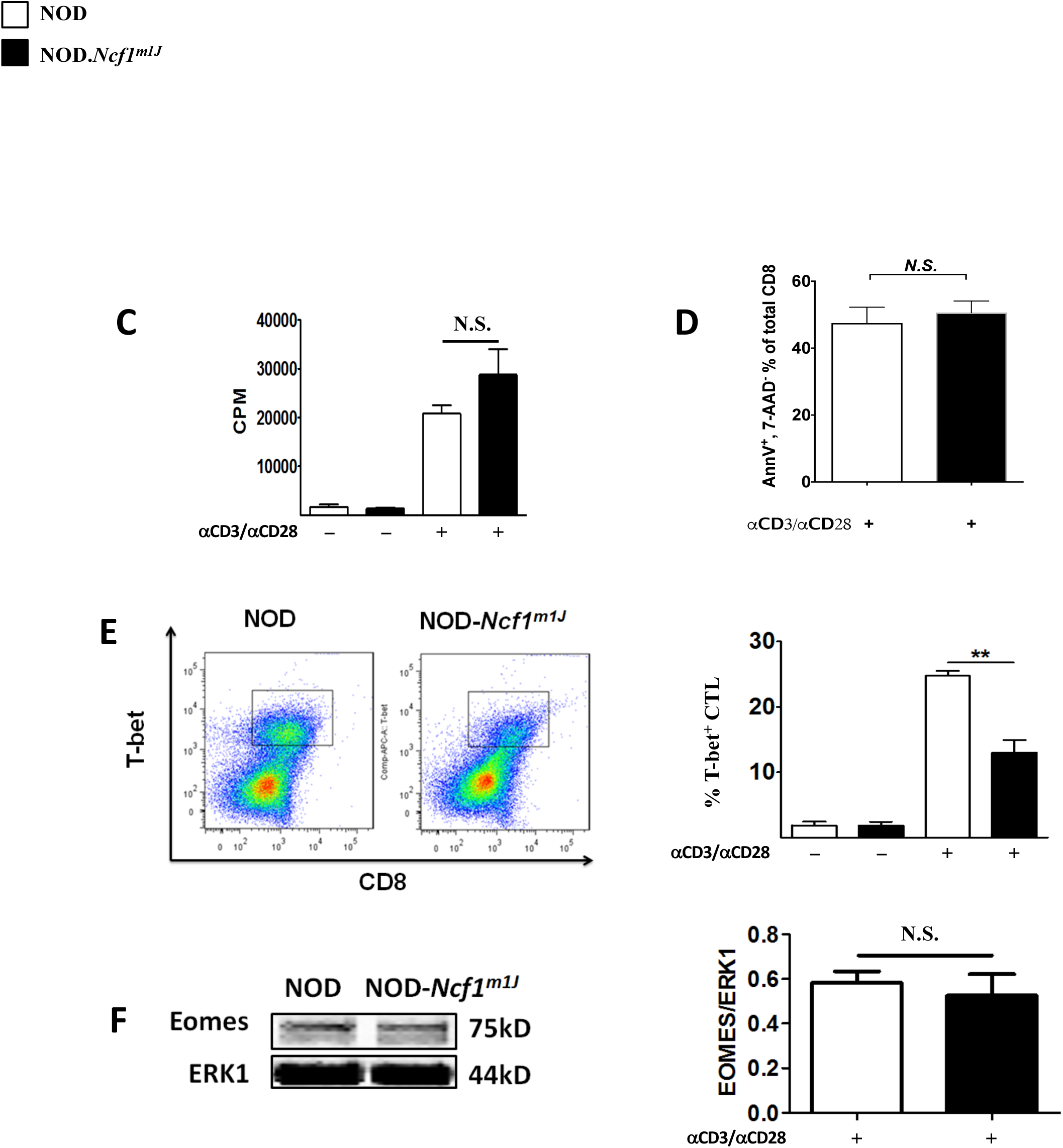
NOD-*Ncf1^m1J^* CD8^+^ T cells exhibit a decrease in pro-inflammatory cytokine and effector molecule production. (A) Representative histogram and the quantitation of intracellular staining in purified CD8^+^ T cells after stimulation of NOD (red histogram or open bar) or NOD-*Ncf1^m1J^* (blue histogram or black bar) with α-CD3/α-CD28 Abs for 72 hours. Gray histogram represents the isotype control. Twelve mice were included in each group. (B) Real-time quantitative PCR of cytokine and effector molecule mRNA in CD8^+^ T cells after α-CD3/α-CD28 stimulation of NOD (open bar) or NOD-*Ncf1^m1J^* (black bar) for 48 hours. PCR was performed with pooled cDNA from three mice and results are representative of three se AI4 splenocytes?independent experiments performed in triplicate. (C) Proliferation was assessed by ^3^H-TdR incorporation using CD8^+^ T cells activated with α-CD3/α-CD28 for 72 hours. D) CTL apoptosis 72 hours after α-CD3/α-CD28 activation. (E) Representative dot-plot and the quantitation of intracellular staining of T-bet in purified CD8^+^ T cells after α-CD3/α-CD28 stimulation for 72 hours. (F) Immunoblot of Eomes in CD8^+^ T cells after stimulation with αCD3/αCD28 for 72 hours. Four mice were included in each group. Data in the bar graphs are represented as mean ± SEM. Statistical analysis used Student’s t test (N.S. *P* > 0.05; * *P* < 0.05; ** *P* < 0.01; *** *P* < 0.001.

In an effort to understand the mechanism behind the selective loss of IFNγ and granzyme B, we sought to investigate the canonical transcription factor of the Tc1 cell lineage, Tbet (39–41). Intracellular staining showed that T-bet levels were decreased in activated NOD-*Ncf1^m1J^* CD8^+^ T cells compared to wild type controls, (Figure 1E). The other transcriptional factor related to IFNγ and granzyme B production, Eomesodermin (Eomes) (39), was present at equal levels in activated NOD and NOD-*Ncf1^m1J^* CD8^+^ T cells (Figure 1F). These results suggest that ROS potentiates T-bet expression to promote CD8^+^ T cell effector function.

### The NOX2 Inhibitor, Apocynin, Suppresses the Production of Effector Molecules in Mouse CD8^+^ T Cells

Production of ROS by NOX2 can be suppressed by apocynin, a specific inhibitor of p47^phox^, the gene product of *Ncf1* (42). Apocynin did not alter CD8^+^ T cell viability (Supplemental Fig 2A) but did inhibit IFNγ production in a dose dependent manner (Supplemental Fig 2B), with 200µM exhibiting maximal effects. To confirm the role of NOX2 in CTL effector function, splenic CD8^+^ T cells were stimulated for 72 hrs with plate-bound α-CD3 and α-CD28 in the presence or absence of 200μM apocynin. Consistent with the observations in NOD-*Ncf1^m1J^*, apocynin dramatically suppressed production of granzyme B, IFNγ, and TNFα protein levels (Figure 2A) and corresponding mRNA transcripts (Figure 2B). Apocynin treated NOD CD8^+^ T cells showed a 30% reduction in intracellular T-bet after α-CD3 and α-CD28 stimulation (Figure 2C). which was similar to results with NOD-*Ncf1^m1J^* CD8^+^ T cells (Figure 1E), apocynin had no impact on proliferation (Figure 2D).

**Figure 2.**
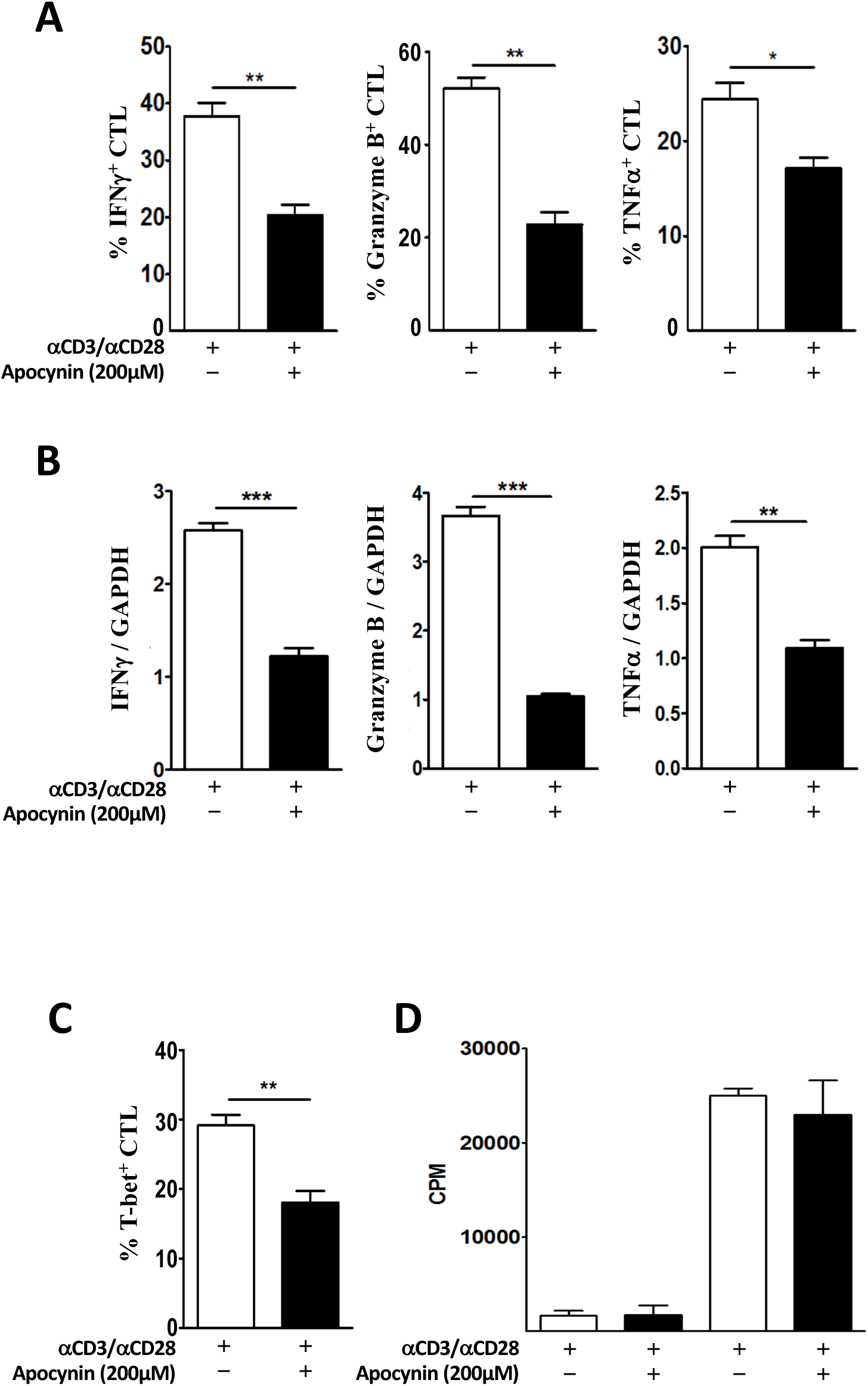
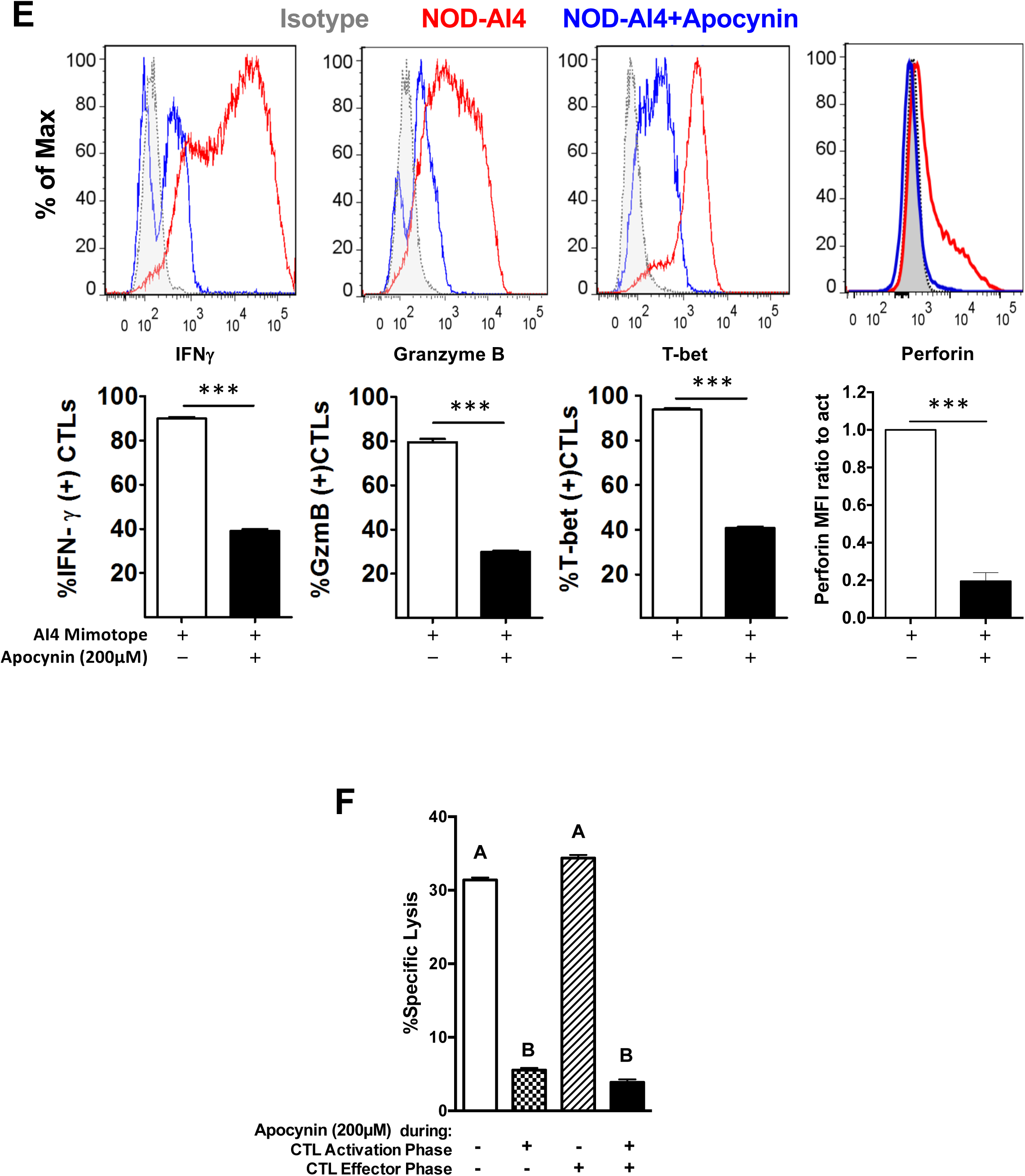
NOX2 is essential for effector function and cytolytic activity of CTL. A-D: CD8^+^ T cells from NOD spleens were activated with plate-bound α-CD3/ α-CD28 with or without the presence of apocynin 200mM. (A) Quantitation of intracellular staining for IFNγ, Granzyme B and TNFα. (B) Real-time quantitative PCR for IFNγ, Granzyme B and TNFα. cDNA was pooled from at least three mice and results represent three independent experiments done in triplicate. (C) Quantitation of intracellular staining of T-bet in NOD CD8^+^ T cells. (D) Proliferation of NOD CD8^+^ T cells. (E) Intracellular staining and quantitative analysis of IFNγ, granzyme B, T-bet, and Perforin expression in autoreactive, monoclonal AI4 CTL after stimulation by specific mimotope for 72 hours either without (open bars) or with (black bars) apocynin (200μM). Gated on Live/Dead_NIR^-^CD3^+^CD8^+.^ (F) AI4-induced CML is significantly reduced when NOX2 was inhibited during T cell activation but not during the effector phase. AI4 T cells were activated by antigen with (CTL Activation phase (+)) or without (CTL Activation phase (−)) apocynin. ^51^Cr labeled NIT-1 NOD-derived beta cells were seeded in 96-well culture plates and cultured with preactivated AI4 cells at an E:T ratio of 20:1 for 16 h, with apocynin (Effector phase (+)) or without apocynin (Effector phase (−)). Data in the bar graphs are represented as mean ± SEM. Statistical analysis used Student’s t test (* *P* < 0.05; ** *P* < 0.01; *** *P* < 0.001). Columns with different letters are statistically different at a level ≤0.05.

To confirm that NOX2 activity during CTL activation was not restricted to the autoimmune-prone NOD mouse, purified splenic CD8^+^ T cells from C57BL/6 (B6) mice were stimulated with α-CD3 and α-CD28 with or without apocynin. T-bet, IFNγ, and granzyme B were assessed by intracellular staining. The percentage of cells positive for the transcription factor T-bet (40.9 ± 2 (B6) vs. 18.8 ± 2.1 (B6+apocynin); p<0.0001) as well as IFNγ (49.2 ± 4.4 (B6) vs. 28.9 ± 1.9 (B6+apocynin); p=0.0027) and granzyme B (50.3 ± 6.6 (B6) vs. 23.2 ± 3.7 (B6+apocynin); p=0.0027) were all reduced in B6 CTL when activated in the presence of apocynin. This phenotype was similar to what was observed in NOD CD8^+^ T cells (Figures 1 and 2). Similarly, B6 cells treated with and without apocynin had comparable upregulation of CD69 and similar levels of proliferation.

### Inhibition of NOX2 Blocks Cytolytic Capacity During Antigen-specific Activation of CTLs

To determine how the loss of NOX2 derived ROS production impacts antigen-specific CTL activation, we used CD8^+^ T cells from the highly pathogenic AI4 TCR-transgenic NOD mouse (NOD.AI4α/β-*Rag1^-/-^*) (35). Upon antigen-specific activation of AI4 cells a stark reduction in the intracellular staining of IFNγ, granzyme B, and T-bet, and perforin occurred when NOX2 was inhibited by apocynin (Figure 2E).

An intact *Ncf1* in CD8^+^ T cells was critical for efficient adoptive transfer of T1D (8) and the loss of NOX2 activity resulted in reduced CD8^+^ T cell cytokine and effector molecule production (Figures 1-2). Thus, we hypothesized that NOX2 activity is essential for cytolytic activity of CD8^+^ T cells. AI4 effector cells were activated by the specific mimotope for 72 hours (activation phase) prior to use in the cell-mediated lymphocytoxicity (CML) assay (effector phase) using the NOD-derived NIT-1 pancreatic β cell line as a target (35). As expected, AI4 cells that were not exposed to apocynin during either the activation or effector phases effectively lysed NIT-1 cells (Figure 2F, 1^st^ bar). β cell killing was prevented when AI4 T cells were treated with apocynin during only the activation phase (Figure 2F, 2^nd^ Bar). In stark contrast, the addition of apocynin during only the CML assay did not affect cytotoxic capability (Figure 2F, 3^rd^ bar). Finally, apocynin treatment of AI4 T cells during both phases significantly reduced lysis. NOX2 is vital for diabetogenic T cell mediated β cell killing and oxidants produced by this enzyme are important during activation/differentiation and likely involved in proximal TCR signaling.

### NOX2 Regulates T-bet, IFNγ, Granzyme B and Effector Function of Human CD8^+^ T Cells

Similar to mouse CD8^+^ T cells, apocynin reduced IFNγ production from human CD8^+^ T cells in a dose dependent manner with 200µM apocynin providing maximal effects (Supplemental Figure 2C). The inhibition of NOX2 in human CD8^+^ T cells with 200µM apocynin led to a significant increase in proliferation (Figure 3A). Whereas proliferation was enhanced in the presence of apocynin, inhibition of NOX2 by apocynin during α-CD3 and α-CD28 activation blunted production of IFNγ, granzyme B, and T-bet (Figures 3B-3D), similar to results with CD8^+^ T cells from NOD and B6 mice. An *in vitro* model of antigen-specific target cell killing was used to determine the impact of NOX2 inhibition on CD8^+^ cytolytic function (Figure 3E). When apocynin (200µM) was absent from both the activation and effector phases, the CTL efficiently lysed the BL5 cells (Figure 3E, open bar). However, when apocynin was added during the activation phase, lysis of BL5 cells fell to less than 10% (Figure 3E, checkered bar). Inhibition of NOX2 during only the effector phase led to a mild reduction in lysis (Figure 3E, hatched bar). Apocynin during both the activation and effector phases resulted in an almost complete inhibition of lysis (Figure 3E, filled bar), comparable to that observed when NOX2 was inhibited during the activation phase alone. These results are similar to the observations using mouse CD8^+^ T cells, where inhibition of NOX2 during CTL activation led to a reduction in T-bet and effector cytokines/molecules with almost complete ablation of cytolytic activity without blunting proliferation. Therefore, NOX2 activity is essential for human and mouse CTL effector function.

**Figure 3.**
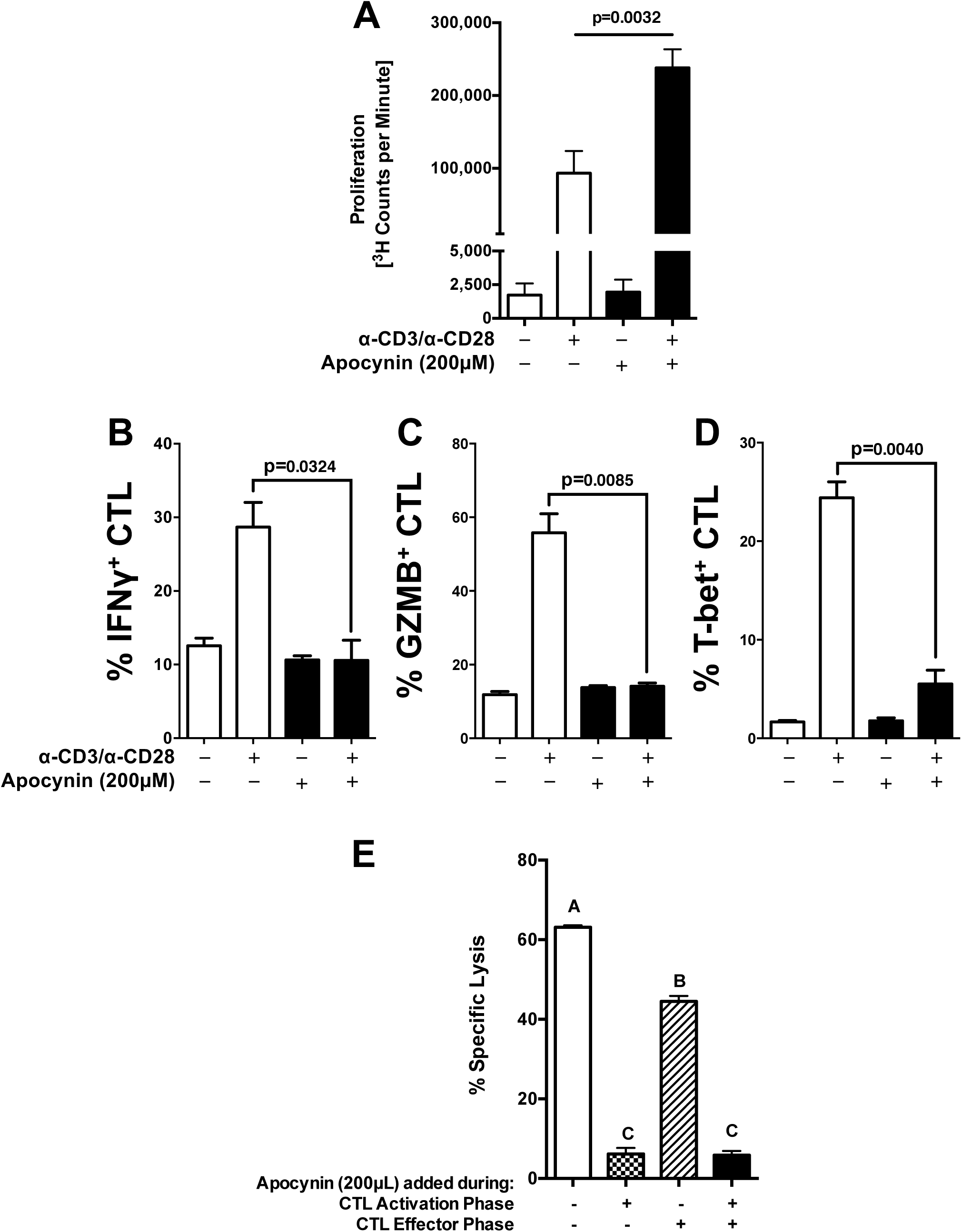
NOX2 is essential for effector function of human CD8^+^ T cells. Purified naïve CD8^+^ T cells from healthy volunteers were activated with αCD3 and αCD28 in the presence or absence of 200µM apocynin. (A) Proliferation was assessed by ^3^H-TdR incorporation at 72 hours. Quantitation of intracellular staining for (B) IFNγ, (C) Granzyme B, and (D) T-bet in purified CD8^+^ T cells (5 × 10^5^ cells) after αCD3 and αCD28 stimulation for 72 hours with or without 200μM apocynin. (E) Purified naïve human CD8^+^ T cells were activated in the presence or absence of 200µM Apocynin for 48 hours, transduced with a lentivirus encoding an IGRP-TCR, and then expanded for an additional 7 days in IL-2 (20U/mL) with αCD3/αCD28 Dynabeads. Apocynin (200µM) was present in specific cultures during the entire CTL activation phase (CTL Activation Phase +). Additionally, CTL were activated in the absence of apocynin (CTL Activation Phase -). The CTL effector phase was performed by co-culturing IGRP reactive CTL at an E:T ratio of 25:1, with 200µM apocynin (CTL Effector Phase +) or without (CTL Effector Phase -). Data represented as mean ± SEM. Statistical analysis used Student’s t test (*** *P* < 0.001). In E bars with different letters are statistically different at a level p<0.05.

### H_2_O_2_, but Not Superoxide, Promotes CD8^+^ T Cell Function

NOX2 produces superoxide as a direct product(9). *In vivo*, superoxide is dismutated to H_2_O_2_ by superoxide dismutase (SOD) family members. H_2_O_2_ can be subsequently catalyzed into H_2_O by catalase (43, 44). Superoxide can travel through the membrane via Chloride Voltage-Gated Channel 3 (ClC-3) and H_2_O_2_ crosses the membrane into the cytosol via aquaporin or diffusion (6, 45). The type of oxidant that modulates the effect of NOX2 in CD8^+^ T cells was examined. Purified NOD CD8^+^ T cells were treated with or without SOD1 and stimulated with α-CD3 and α-CD28 conjugated beads. SOD1 had little impact on the synthesis or secretion of IFNγ (Figures 4A-4B). In addition, SOD1 had no effect on synthesis of granzyme B or TNFα (Figure 4A). Accordingly, SOD1 treatment had no effect on T-bet (Figure 4C).

**Figure 4.**
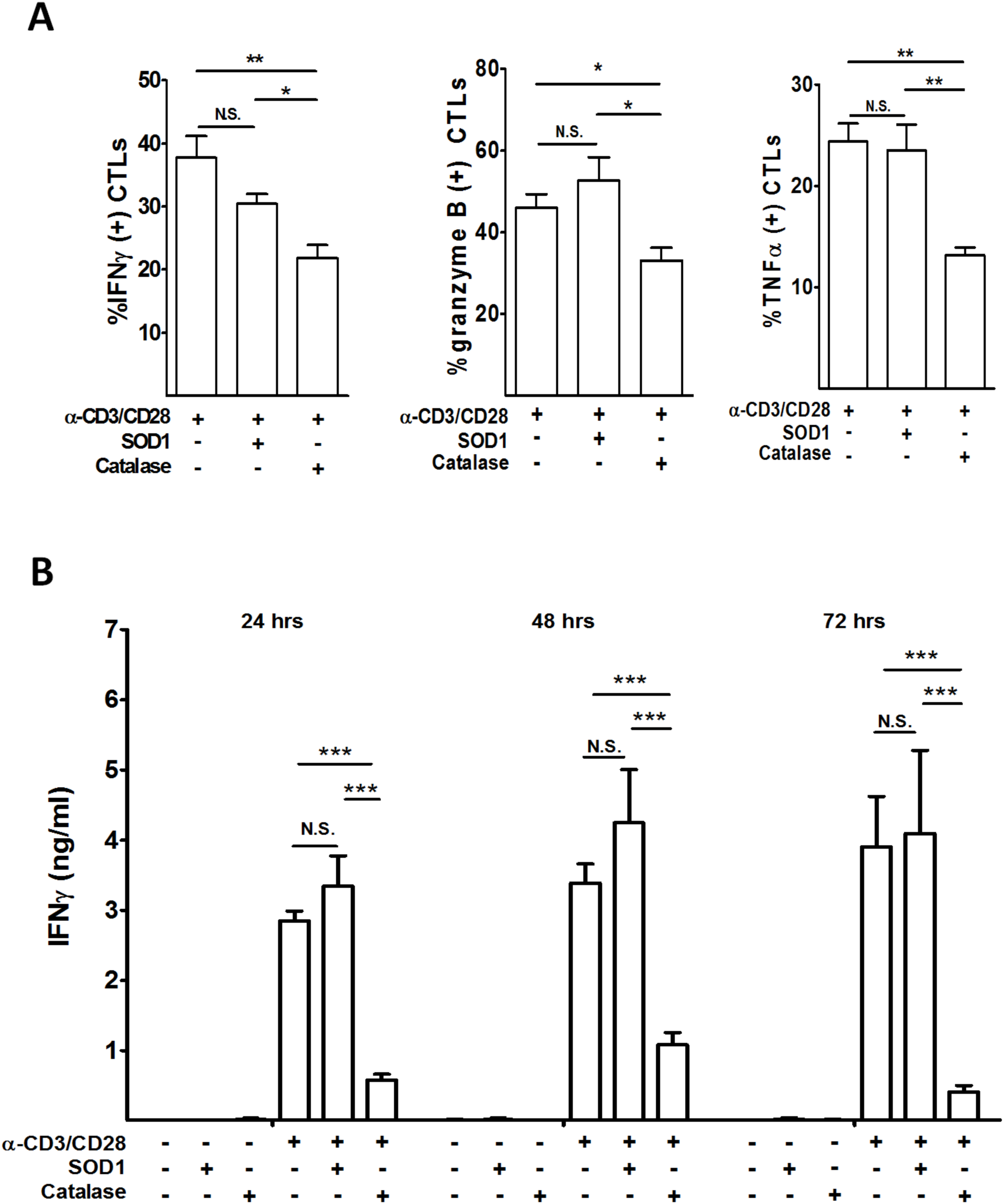
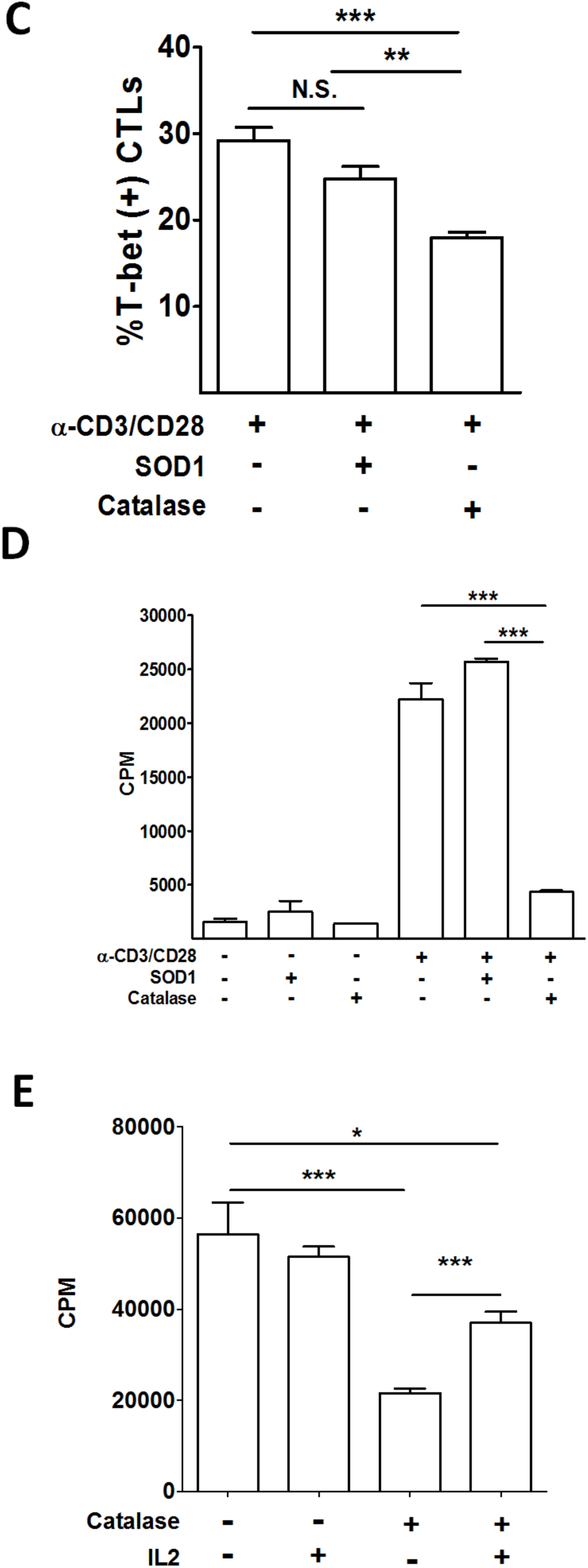
Hydrogen Peroxide scavenging significantly blunts CTL effector function after αCD3/αCD28 stimulation. CTL were stimulated by αCD3/αCD28 for 3 days in the presence or absence of catalase or SOD1. (A) Intracellular levels of IFNγ, granzyme B and TNFα were assessed. (B) IFNγ was measured by ELISA on days 1, 2 and 3. (C) Intracellular staining for T-bet in CTL. (D) CTL proliferation was assessed by [^3^H]TdR incorporation. (E) CTL proliferation was measured at 72 hours after αCD3/αCD28 stimulation in the presence of catalase and IL2. Results are presented as mean ± SEM. Each measure was compiled from four independent observations performed in triplicate. (* *P* <0.05; ** *P* <0.01; *** *P* <0.001).

To examine the effect of H_2_O_2_, purified NOD CD8^+^ T cells were treated with or without catalase and stimulated with α-CD3 and α-CD28. Catalase treatment significantly suppressed production of IFNγ, granzyme B and TNFα (Figures 4A and 4B). Furthermore, T-bet expression after α-CD3 and α-CD28 stimulation also was significantly reduced following catalase treatment (Figure 4C). Therefore, H_2_O_2_ transmits the redox signal to promote T-bet expression and effector function in CD8^+^ T cells. Interestingly, unlike NOD-*Ncf1^m1J^* or apocynin-treated cells, CD8^+^ T cell proliferation was dramatically suppressed by scavenging H_2_O_2_ (Figure 4D). This is consistent with role for a NOX2-independent ROS source, such as mitochondrial ROS, in IL-2 production and CD8^+^ T cell proliferation (46). Indeed, addition of IL-2 to the catalase treated CTL partially rescued cell proliferation (Figure 4E).

To determine if the defective effector functions associated with loss of NOX2 activity persist when proximal signaling events are bypassed, NOD and NOD-*Ncf1^m1J^* CD8^+^ T cells were activated with PMA and ionomycin (47). When NOD-*Ncf1^m1J^* CD8^+^ T cells were activated by PMA and ionomycin, protein levels of IFNγ, granzyme B and T-bet were equal to NOX2-intact NOD CD8^+^ T cells (Supplemental Figure 3A). Similarly, catalase treatment did not impact CD8^+^ T cells activated with PMA/ionomycin (Supplemental Figure 3B). Therefore, NOX2-derived H_2_O_2_ acts proximally but not distally in TCR signaling.

### NOX2 Activity Impacts TCR signaling at the TSC1/2 Complex to Promote mTORc1 Activity

Proximal TCR signaling events include activation of AKT by PI3K and subsequent repression of the TSC1/2 complex (38). The inactivation of TSC1/2 results in elevated RheB-GTP levels that can promote mammalian target of rapamycin complex 1 (mTORc1) activity leading to downstream effects on T-bet, IFNγ, and granzyme B (5). The AKT pathway was not altered, as lysates of α-CD3 and α-CD28-stimulated NOD and NOD-*Ncf^m1J^* CTLs displayed comparable p-GSK levels (Figure 5A). However, differences in TSC1/2 activity were observed, as RheB-GTP increased ∼2-fold after activation in NOD CTL, whereas an increase in RheB-GTP was not observed in NOD-*Ncf1^m1J^* CTL (Figure 5B).

**Figure 5.**
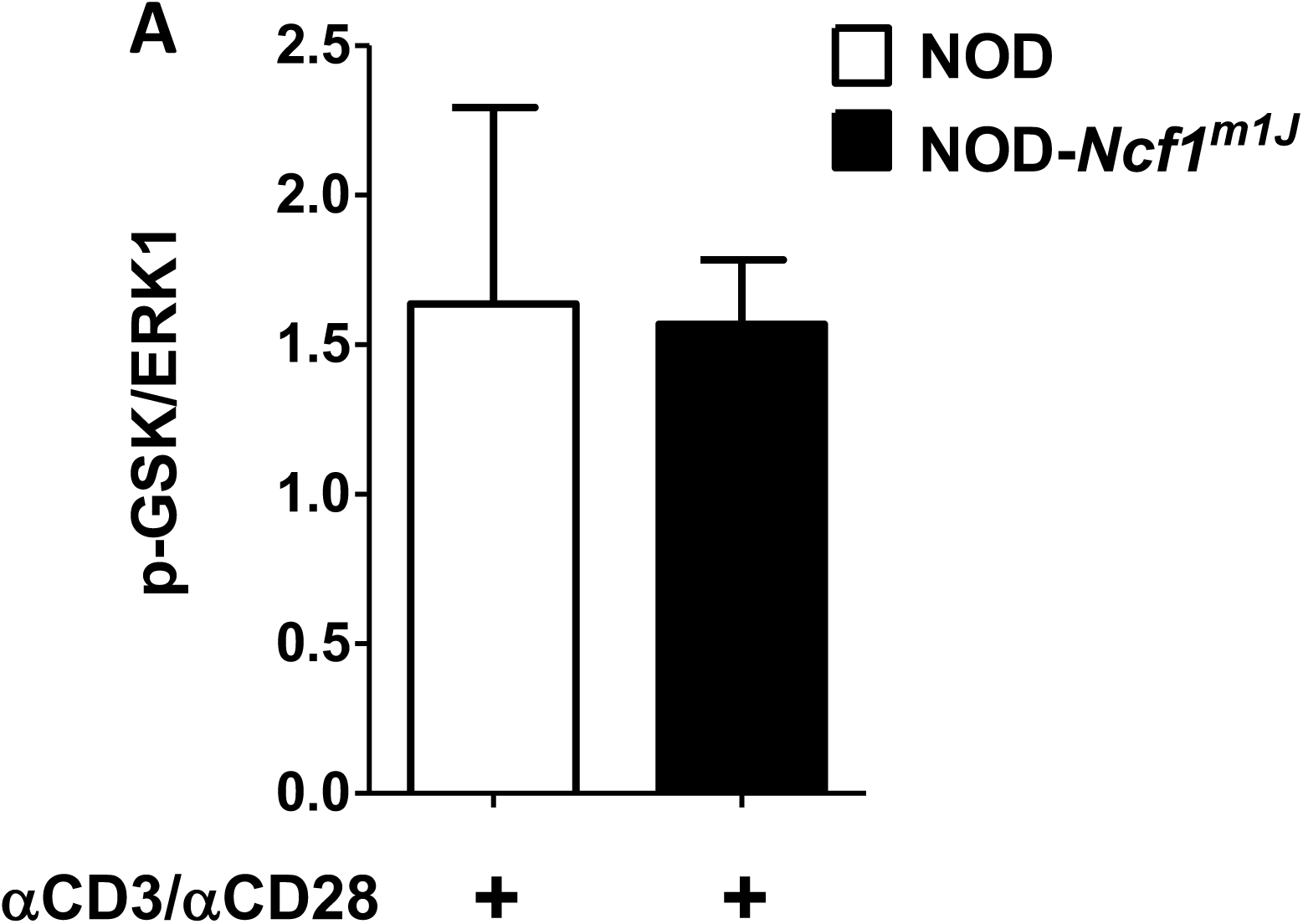
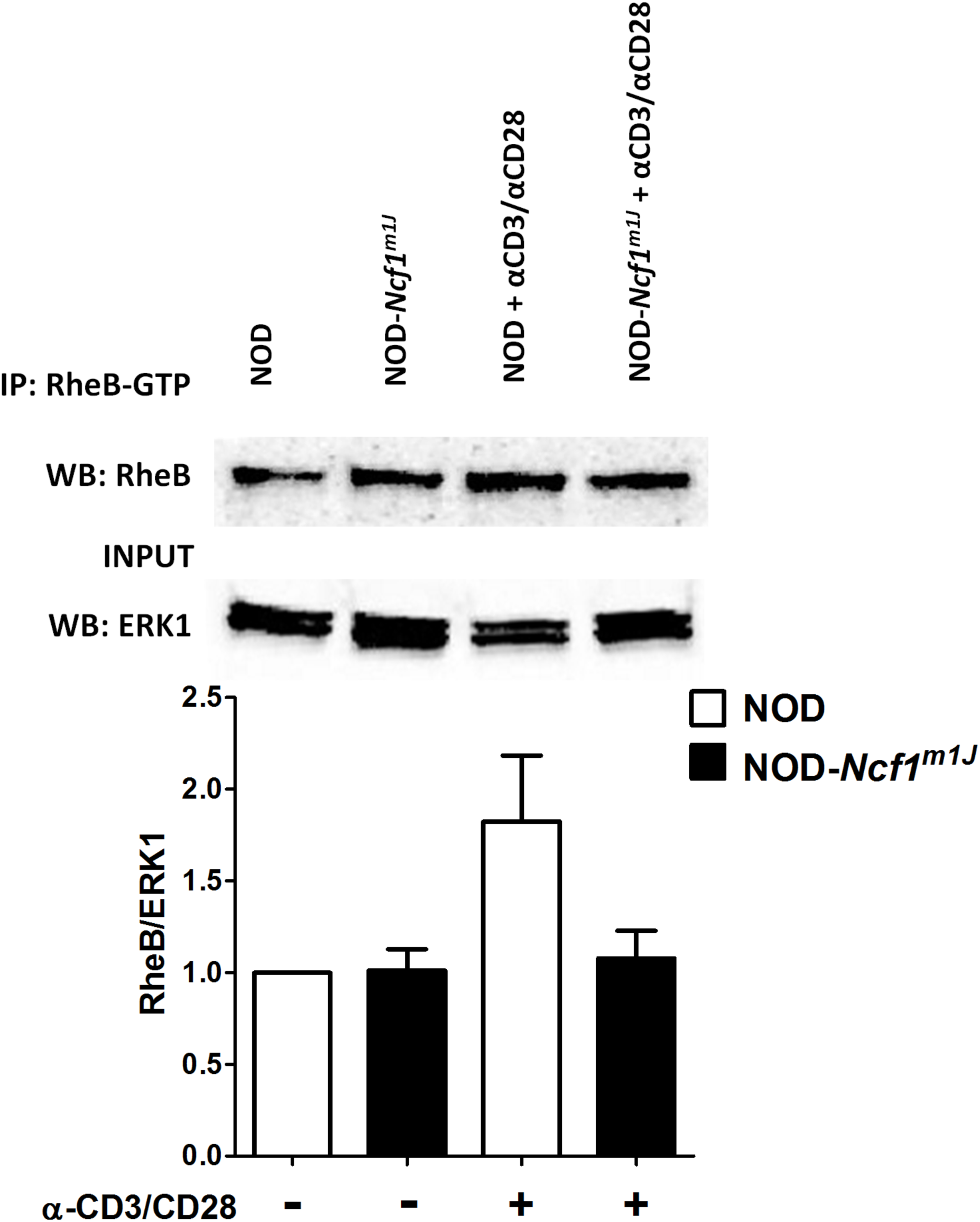
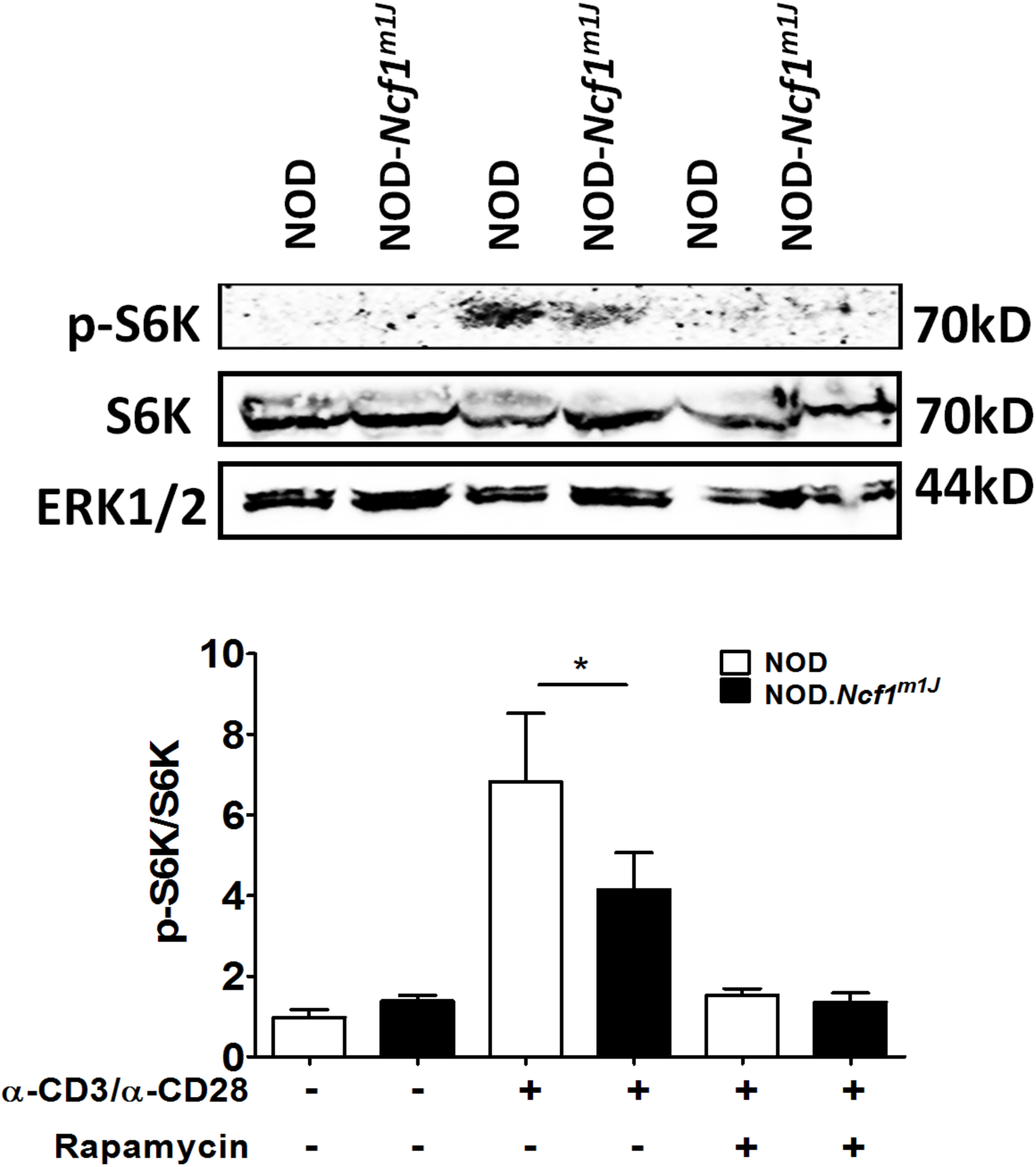
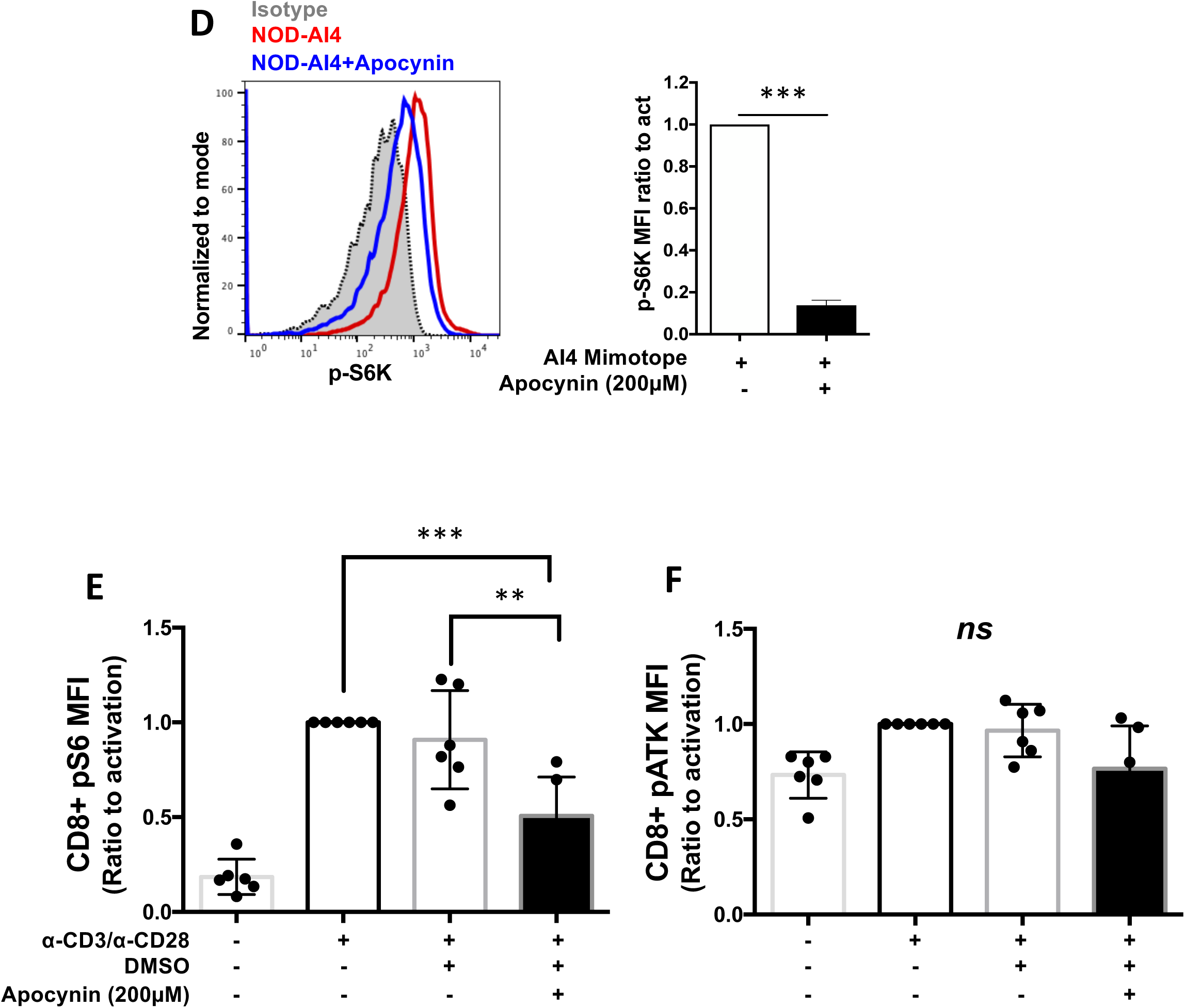
NOX2 activity is required for TSC1/2 repression and downstream mTORc1 activation during CTL activation. NOD or NOD-*Ncf1^m1J^* CD8^+^ T cells were activated by α-CD3/α-CD28 for 30 minutes and subjected to immunoprecipitation of p-GSK and RheB-GTP. (A) Cellular p-GSK levels, a marker of Akt kinase activity, were determined relative to the input control Erk1/2. (B) Cellular level of RheB-GTP was determined by calculating the ratio of immunoprecipitated RheB-GTP and input control Erk1/2. Untreated NOD CTL was used as the reference group for normalization between different tests. (C) Immunoblot showing a compromised mTOR complex activity in CTL from NOD-*Ncf1^m1J^*. NOD or NOD-*Ncf1^m1J^*. CD8^+^ T cells were activated by α-CD3/α-CD28 for 30 minutes with the presence or absence of 20ng/mL rapamycin. Phosphorylation of S6K was measured by the ratio of phosphorylated S6K and total S6K. Untreated NOD CTL was used as the reference group for normalization among multiple tests. (D) Intracellular staining and quantitative analysis of pS6K in autoreactive, monoclonal AI4 CTL after stimulation by specific mimotope for 72 hours either without (open bars) or with (black bars) apocynin (200μM). Gated on Live/Dead_NIR^-^CD3^+^CD8^+^. (E-F) Human PBMC from healthy donors (n=6) with or without pretreatment of apocynin (400 µM) were activated with anti-CD3, anti-CD28 for 10 min. Surface markers and intracellular p-S6 and p-AKT were stained with fluorescent antibodies and detected using flowcytometer. Showing are p-S6 (top panel) and p-AKT (Lower panel) MFI ratio to activated, gated on Live/Dead-Yellow^-^CD3^+^CD8^+^ T cells. Results are compared using Student’s t test or One-way ANOVA Multicomparisons (* *P* < 0.05; ** *P* <0.01; *** *P* <0.001).

mTORc1 is essential for CD8^+^ T cell effector function through promoting T-bet expression (5). In fibroblasts, mTORc1 activity can be promoted by oxidants through a process that is dependent on oxidation and repression of TSC1/2 function (48). We therefore hypothesized that NOX2-derived ROS may enhance CTL effector function by promoting mTORc1, an upstream mediator of T-bet. mTORc1 phosphorylates S6K at T389 (p-S6K) and can be used as an indicator of mTORc1 activity (48, 49). Purified CTLs from NOD mice exhibited a significant increase in p-S6K upon TCR activation, indicating sufficient mTORc1 activity after stimulation. Meanwhile, S6K phosphorylation was significantly lower in activated NOD-*Ncf1^m1J^* CD8^+^ T cells (Figure 5C). This was further confirmed through inhibition of NOX2 by apocynin leading to lower p-S6K in antigen-specific activation of mouse AI4 CD8^+^ T cells (Figure 5D). In addition, we were able to show the same effect of apocynin in human CD8^+^ T cells. Lack of NOX2 function inhibited activation-induced P-S6 (Figure 5E) but not p-AKT (Figure 5F) in human CD8^+^ T cells. These data support the hypothesis that a functional NOX2 inactivates TSC1/2 resulting in downstream TCR-dependent RHEB-GTP and mTORc1 activation in both mouse and human CD8^+^ T cells.

To confirm that mTORc1 was required for CD8^+^ T cell effector function, we treated CTL with the mTORc1 inhibitor rapamycin and activated the cells with α-CD3 and α-CD28. Inhibition of mTORc1 dramatically reduced IFNγ, granzyme B, and TNFα production in CD8^+^ T cells (Figure 6A). Suppression was observed as early as 24 hours after stimulation, indicating that inhibition of mTORc1 interrupts early events during CTL activation (Figure 6B). Consistent with a previous report (5), the compromised effector function results from a reduction in T-bet production after activation of rapamycin treated CD8^+^ T cells (Figure 6C).

**Figure 6.**
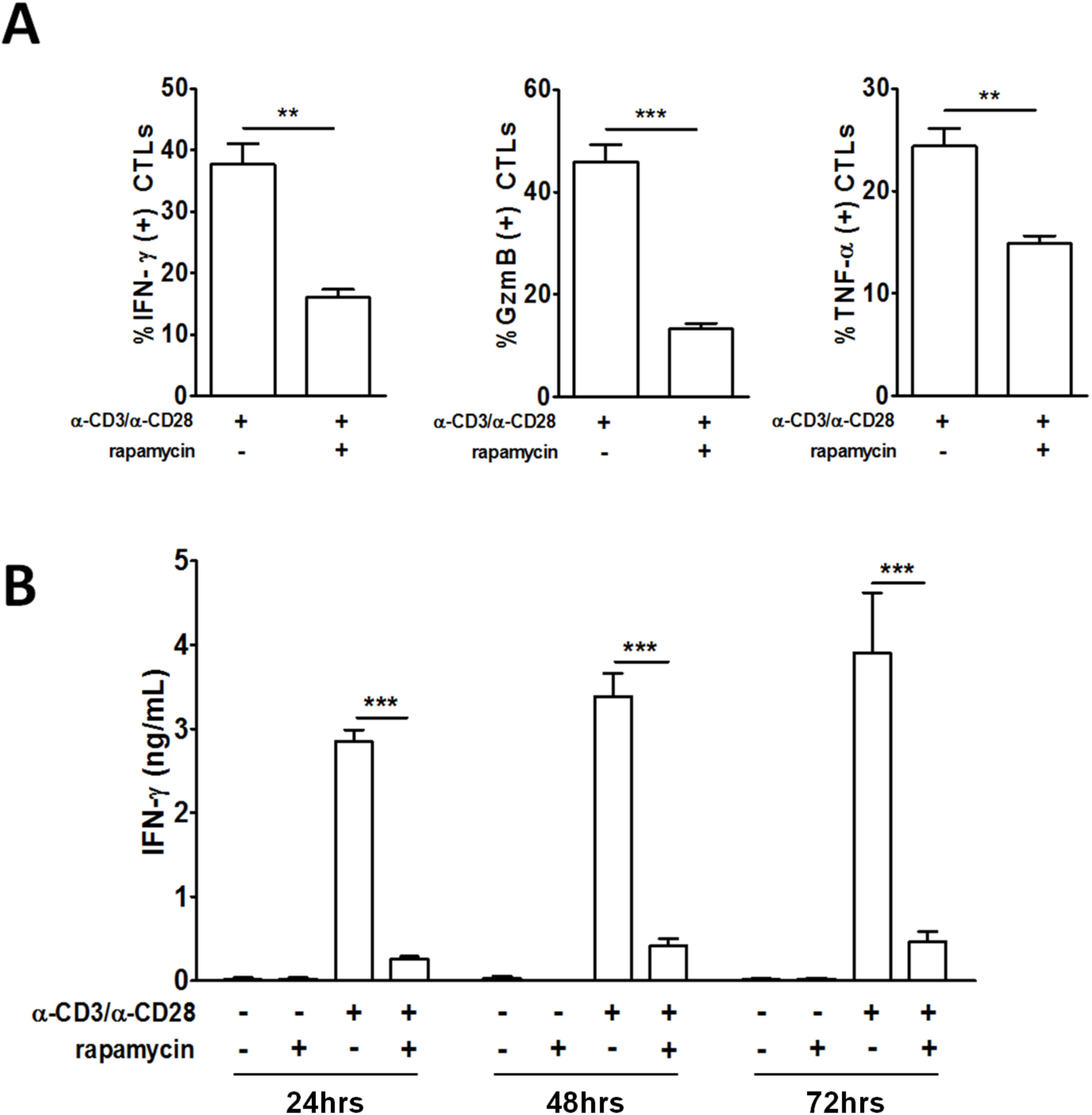
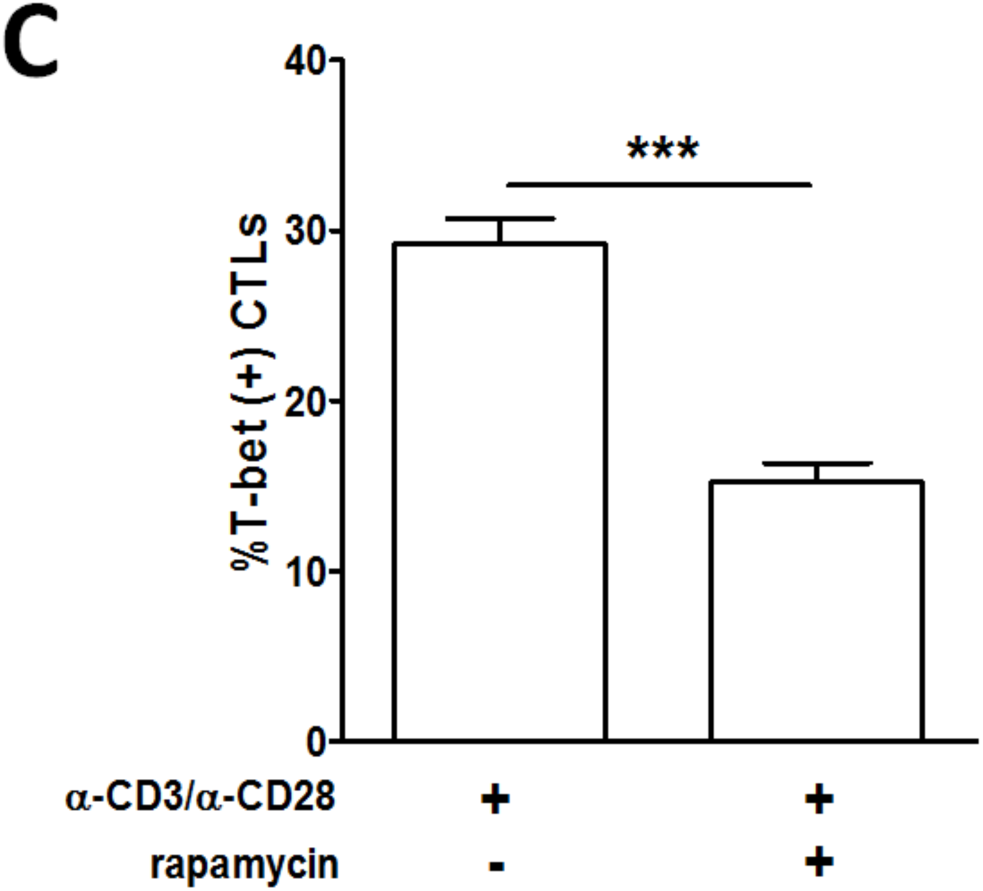
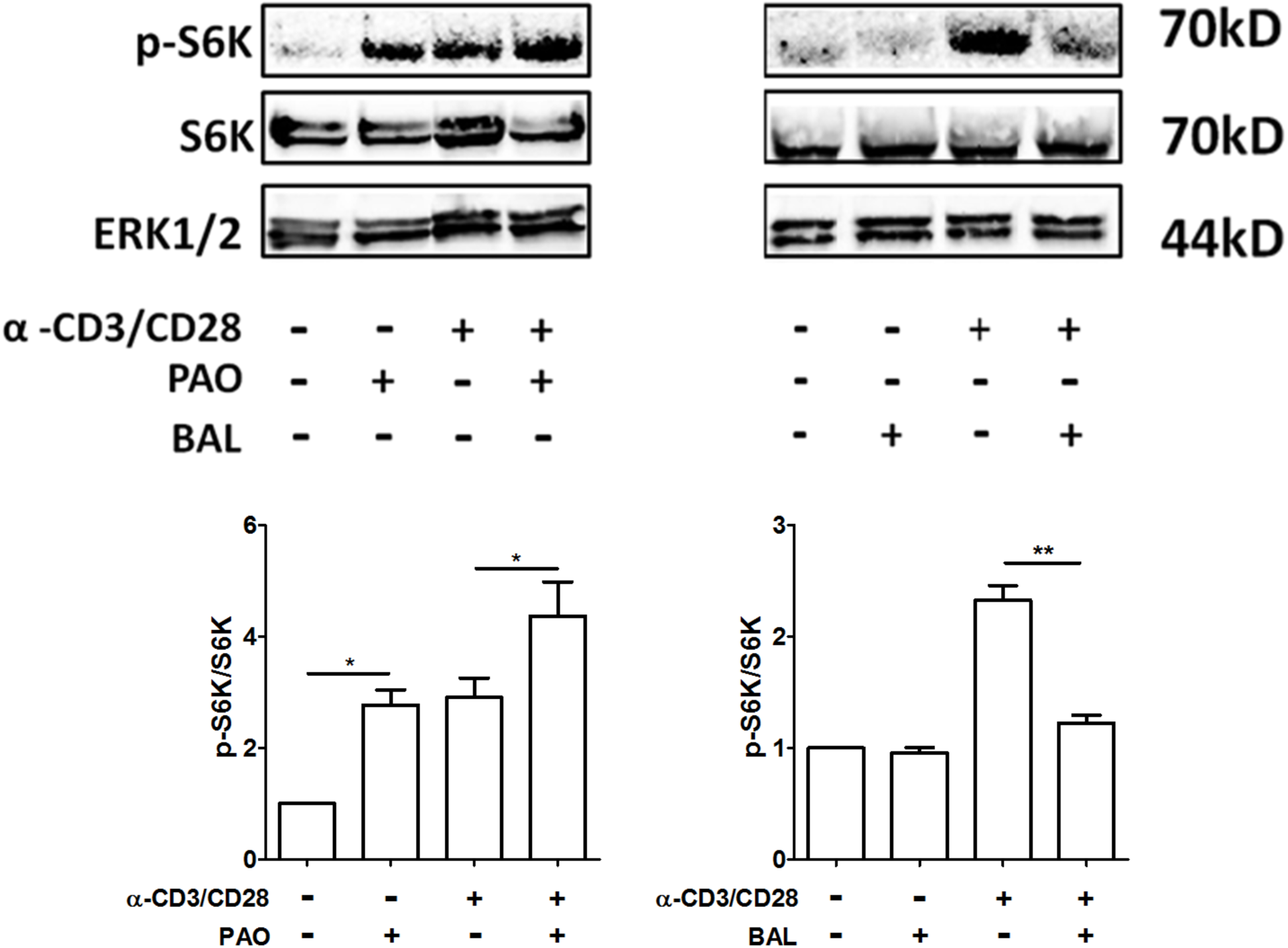
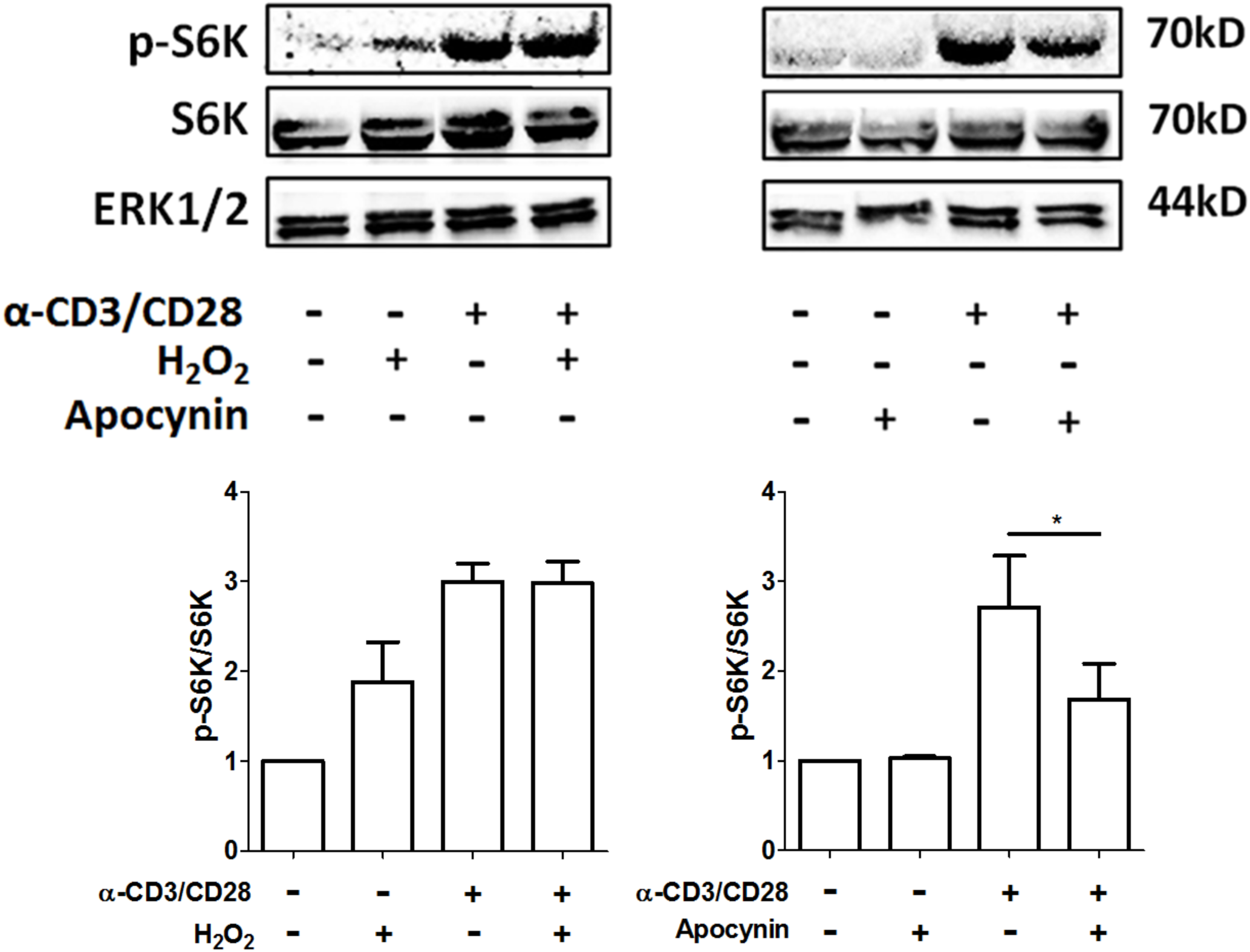
mTORc is required for CD8^+^ T cell effector function through promoting transcriptional activity of T-bet through a redox-regulated regulated mechanism. (A) Quantitation of intracellular staining of the IFNγ, granzyme B, and TNFα in purified NOD CD8^+^ T cells stimulated by α-CD3/α-CD28 for 66 hours in the presence or absence of 20ng/mL rapamycin followed by re-stimulation with PMA/ionomycin plus Golgi stop for 6 hours. (B) NOD CD8^+^ T cells (5 × 10^4^) were stimulated by α-CD3/α-CD28 in the presence or absence of 20ng/mL rapamycin. IFNγ was measured by ELISA on day 1, 2 and 3. (C) Intracellular staining for T-bet in CTL after a 72-hour polyclonal stimulation. NOD CD8^+^ T cells were treated with (D) PAO or BAL or (E) H_2_O_2_ or apocynin either with or without activation by α-CD3/α-CD28 for 30 minutes. Phosphorylation of S6K was measured by the ratio of phosphorylated S6K and total SK6. Results are from three independent experiments. Untreated NOD CTL were used as the reference group for normalization among multiple tests. Results are presented as mean ± SEM. Student’s t-test was performed to exclude potential batch effects (N.S. not statistically significant; * *P* < 0.05; ** *P* < 0.01; *** *P* < 0.001)).

### Redox Regulation of mTORc1 Activity in CD8^+^ T cell

The compromised mTORc1 activity in NOD-*Ncf1^m1J^* (Figure 5C) led us to propose that mTORc1 activity in NOD CD8^+^ T cells is redox-dependent. To explore this possibility, the effect of oxidants or antioxidants upon mTORc1 activation was assessed. Phenylarsine oxide (PAO)is a cell permeable reagent that cross-links vicinal thiol groups and thus, was used as a cysteine oxidant. BAL was used as a reducing reagent. According to previous reports, PAO is able to significantly promote the phosphorylation of S6K by mTORc1 in HEK293T cells, while BAL blocks the PAO-induced or endogenous oxidant mediated mTORc1 activation in these cells (48, 50). NOD CD8^+^ T cells were treated with either PAO or BAL in the presence or absence of α-CD3 and α-CD28 for 45 minutes. In the absence of TCR activation, phosphorylation of S6K was increased by PAO (Figure 6D). When CTL were treated with BAL, p-S6K was dramatically decreased to basal levels even in the presence of α-CD3 and α-CD28 stimulation (Figure 6D). This confirmed the positive effect of oxidants on mTORc1 in CD8^+^ T cells. As our data demonstrates that H_2_O_2_ could be the effector molecule of NOX2 to elicit the redox signal in T cell activation, we tested the ability of H_2_O_2_ to promote mTORc1 activity in CD8^+^ T cells. NOD CTL were treated with 10μM H_2_O_2_ and then activated with α-CD3 and α-CD28. Similar to the PAO treatment of resting CD8^+^ T cells, H_2_O_2_ promoted phosphorylation of S6K even in the absence of polyclonal stimulation. However, such a low dose of H_2_O_2_ did not potentiate mTORc1 activity during polyclonal activation of T cells. A plausible explanation was that this resulted from redundant ROS produced in NOX2-intact CTL after TCR engagement (Figure 6E). To confirm NOX2 as the H_2_O_2_ source, CTL were treated with apocynin and S6K phosphorylation was assessed with or without α-CD3 and α-CD28 stimulation. Consistent with NOD-*Ncf1^m1J^* CD8^+^ T cells, when NOX2-intact CTL were activated in the presence of apocynin, p-S6K levels were significantly reduced (Figure 6E). Taken together, these data demonstrate that NOX2-derived H_2_O_2_ by CD8^+^ T cells can mediate TCR signaling through mTORc1 via TSC1/2 as shown in our proposed model (Figure 7).

**Figure 7.**
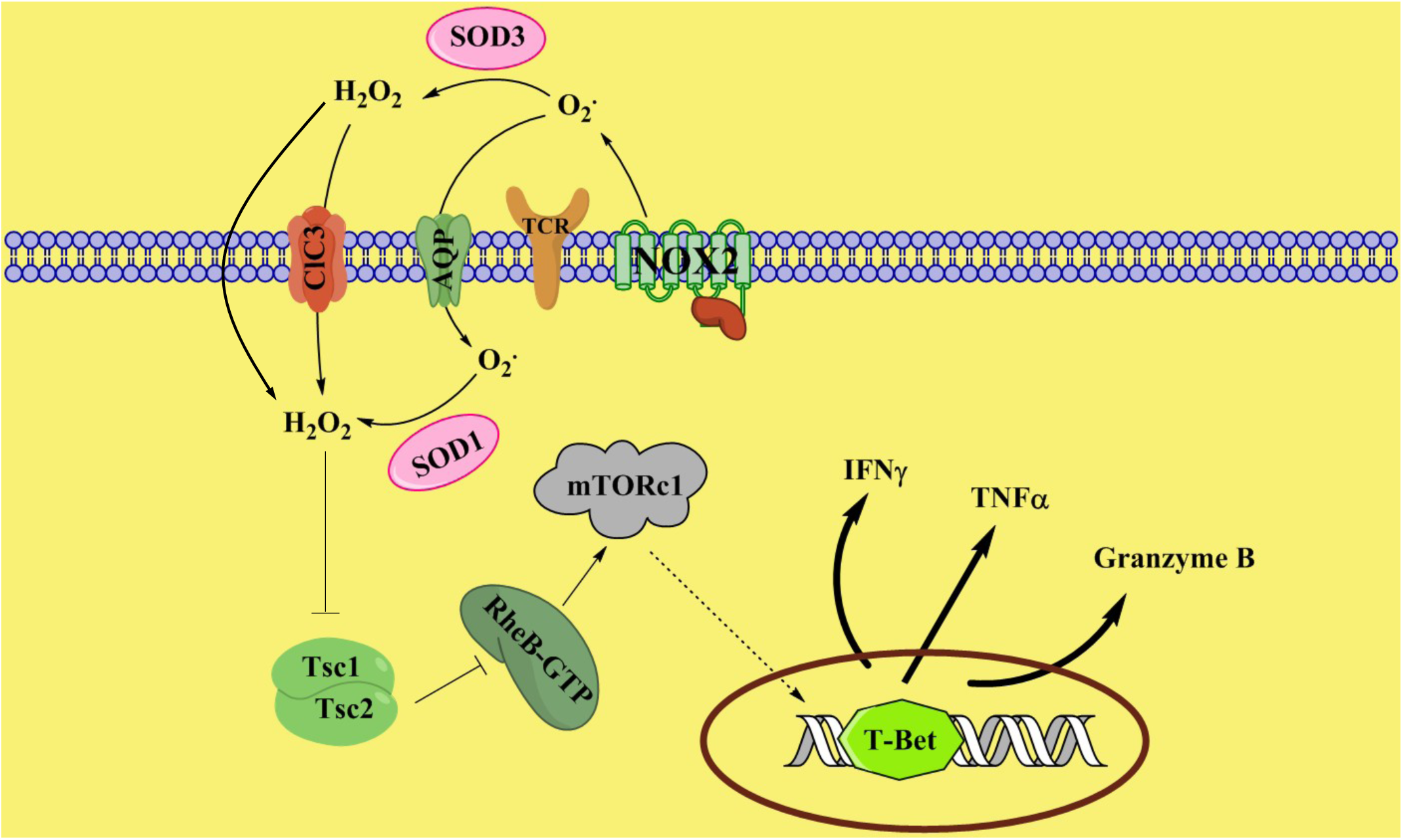
Proposed model of NOX2-mediated redox signaling during activation of CTLs. Upon activation, NOX2 generates superoxide, which is dismutated into hydrogen peroxide, probably by superoxide dismutase. H_2_O_2_ in turn inhibits Tsc1/2. Suppression of Tsc1/2 activity promotes the levels of RheB-GTP, leading to enhanced mTORc1 activity. Up-regulation of mTORc1 activity promotes T-bet expression and thus facilitates CTL effector function. NOX, NADPH oxidase; O_2_^•^, superoxide; H_2_O_2_, hydrogen peroxide; SOD, superoxide dismutase.

## DISCUSSION

During the interaction of naïve T cells with antigen-presenting cells, TCR signaling is coordinated with secondary and tertiary signals to decide T cell fate and differentiation (51). Previous studies have demonstrated that ROS govern processes of T cell activation, including IL-2 production, proliferation, differentiation, and effector function (7, 11, 52–55). In T cells, various ROS generators collaboratively promote cell activation and govern effector phenotypes (46, 54). Here, we have identified that another ROS generator, NOX2, is vital for CD8^+^ T cell activation by promoting effector function and cytotoxicity through the mTORC1/T-bet axis signaling pathway (Figure 7).

T1D is primarily mediated by autoreactive T cell responses. Both clinical and basic science reports have provided evidence that support a role for ROS in the pathogenesis of T1D (56). CD8^+^ T cells are recognized as a major effector cell type in mediating β cell death in T1D (22, 57, 58). Multiple reports suggest that the transcriptional factor T-bet is required in the development of T1D by promoting CTL effector differentiation (59–61). Our previous studies have indicated that NOX2 promotes T_H_1 differentiation via T-bet in NOD CD4^+^ T cells (11) and is required for full effector function of CD8^+^ T cells in T1D using both NOD-*Ncf1^m1J^* as well as adoptive transfer models. Here, we have discovered a novel mechanism where NOX2-derived ROS promotes CTL effector function through enhancing T-bet expression. Stimulated-NOD*-Ncf1^m1J^* CTL showed a reduction in T-bet expression and suppression of CTL effector responses including IFNγ, granzyme B, and TNFα production. Similarly, when treated with the NOX2 inhibitor apocynin, polyclonal activation of CD8 T cells from the NOD and B6 strains showed a decline in T-bet synthesis. These results were extended to human model systems where inhibition of NOX2 during activation prevents T-bet, IFNγ, and granzyme B expression and effector function of both human and mouse CTL.

The treatment of diabetogenic AI4 CTL with apocynin during antigen-specific activation resulted in an ablation of T-bet, IFNγ, and granzyme B as well as β cell-cytolytic capability. In contrast, if NOX2 was only inhibited during the CML assay, the lysis of β cells was equal to untreated AI4 CTL, suggesting that the effect of NOX2 is early during CTL activation and is not required for CTL degranulation. These results were confirmed in a human model effector:target co-culture system. When NOX2 was blocked by apocynin during the activation phase, IGRP-reactive human CTL-avatars could not lyse human β cells. The fact that specific lysis of β cells was not impacted or only minimally affected by addition of apocynin during the effector phase of the CML assay supports the notion that blocking NOX2 activity within the β cell would not provide significant protection of the beta cells from lysis by CTL. The results corroborate our previous studies where NOD-*Ncf1^m1J^* mice were susceptible to adoptive transfer of AI4 CTL (8). These data highlight an important role for NOX2 in promoting CTL effector function during the early events of activation. CTL proliferation was not impacted when NOX2 was inactive due to genetic ablation or pharmacological inhibition, which is consistent with a previous report on the proliferative capacity of T-bet knock-out CTL (60).

NOX2 generates superoxide at the exterior side of the cell membrane (9, 10). Superoxide is thought to be transported across membranes by ClC-3, a member of the ClC voltage-gated chloride (Cl^-^) channel superfamily. Alternatively, it could be converted into H_2_O_2_ outside of CTL that can then diffuse into the cytosol or be transported by aquaporins (45, 62, 63). Scavenging superoxide by treating CTL with SOD1 during polyclonal activation had little effect on proliferation or expression of T-bet, IFNγ, granzyme B or TNFα; however, scavenging H_2_O_2_ resulted in significant decreases of these CTL effector molecules. This indicates H_2_O_2_ instead of superoxide modulates CTL effector functions. CTL proliferation also was dramatically suppressed by catalase treatment, suggesting a NOX2-independent source of ROS was functioning to promote CTL proliferation. As mitochondrial ROS promotes T cell proliferation by enhancing IL-2 production (46), we conclude that in CTL, NOX2 and mitochondrial-derived H_2_O_2_ enhance T cell activation by promoting effector differentiation and proliferation, respectively. Indeed, when IL-2 was added to the cultures during CTL activation with α-CD3/α-CD28 in the presence of catalase, proliferation was restored.

NOX2 promotes T-bet production through modulating mTORc1 activity after α-CD3 and α-CD28 stimulation of naïve CD8^+^ T cells. mTORc1 promotes the production of T-bet over its counterpart Eomesodermin after CTL activation, to regulate the balance between CTL effector and memory differentiation (5). Inhibition of mTORc1 with rapamycin had a dramatic suppressive effect on NOD CTL T-bet production, pro-inflammatory cytokine, and effector molecule expression, but with little impact on cell proliferation. CTLs with *Ncf1* mutation or inhibition were compromised in TCR-dependent mTORc1 activation. Our results provide additional data to add to a more complete picture of mTORc1 activation and induction of differentiation in CD8^+^ T cells (64–68) to include NOX2-derived hydrogen peroxide as an upstream mediator of mTOR.

Hydrogen peroxide derived from NOX2 promotes CTL effector function. Hydrogen peroxide treatment of CTL activated mTORc1; however, when NOX2 produced enough free radicals upon α-CD3 and α-CD28 stimulation, H_2_O_2_ could not further promote mTORc1 activation. Treating NOD CTL with cysteine specific oxidant, PAO, greatly enhanced mTORc1 activity, even without the CD3/CD28 signal. In contrast, the antioxidant BAL inhibited the activation of this complex in CTL, confirming the significance of ROS in mTORc1 activation. mTORc1 activation is involved in multiple cellular pathways and RheB-GTP, the active GTP bound form of RheB, is a major activator of mTORc1 by direct interaction (69, 70). In many cell types, RheB-GTP is negatively regulated by TSC1/2 by transforming RheB-GTP into Rheb-GDP, the inactive form (71). Our data demonstrate that in the absence of NOX2-derived ROS, RheB-GTP levels are not suppressed indicating compromised TSC1/2 activity (Figure 5B). Akt function was not impaired in the absence of NOX2, demonstrating that NOX2 regulates TSC1/2 activity in CTL in an Akt-independent manner, as previously described using immortalized cell lines (48).

In summary, this study provides a novel mechanism whereby oxidants regulate CD8^+^ T cell effector function. NOX2-derived H_2_O_2_ enhances Rheb activity and thus promotes mTORc1 function during CTL activation, leading to an activation of T-bet as well as downstream CTL effector function and cytolytic activity. The data reported here do not support a global pro-inflammatory dysfunction in NOX-deficient animals. *Ncf1* mutations previously have been associated with increased severity of experimental allergic encephalomyelitis and collagen-induced arthritis (11, 72) yet resistance to T1D (8, 11, 25). Due to the essential role of CD8^+^ T cells in T1D NOX2-derived ROS play CTL extrinsic and intrinsic roles in pathogenesis. NOX2 promotes dendritic cell antigen cross presentation and activation of naive CTL in T1D (extrinsic (25)) and here we provide evidence for a CTL intrinsic role of NOX2 in regulating signal transduction leading to production of effector molecules and cytolytic function (Figures 1-4 and 7). Our data provide new insights into potential targets for therapeutic strategies in organ transplantation or CTL-mediated autoimmune disease, such as T1D. In addition, due to the essential role of ROS in immune cell signal transduction, we predict that non-targeted antioxidant therapy or supplements might alter the thresholds for CTL effector function during immune responses.

## ACKNOWLEDGEMENTS

This work was supported by research grants from the JDRF, the National Institutes of Health R01 DK074656 (C.E.M.), UC4 DK104194 (C.E.M.), P01 AI042288 (C.E.M./T.M.B), F30 DK105788 (B.N.N.), Horizon Therapeutics (J.W.L.) and the Sebastian Family Endowment for Diabetes Research. The authors declare that no conflicts of interest exist.

## Author Contributions

JC researched the data and wrote the manuscript, CL researched the data and wrote the manuscript, AVC researched the data and reviewed/edited the manuscript, BNN researched the data and reviewed/edited the manuscript, TMB provided key resources, contributed to discussion, and reviewed/edited the manuscript, YX researched the data and reviewed/edited the manuscript, NM researched the data and reviewed/edited the manuscript, CS researched the data and reviewed/edited the manuscript, WHR contributed to discussion and reviewed/edited the manuscript, HMT conceived of study components and reviewed/edited the manuscript, JWL researched the data and reviewed/edited the manuscript, and CEM conceived of the study, oversaw all aspects of the research, and wrote the manuscript.

## Notes

### Competing Interest Statement

The authors have declared no competing interest.

### Summary of Updates

In the original submission the x axis label for one group in Figure 3 was mislabeled. This has been corrected in the resubmission.

