## Supplementary material for "NADPH Oxidase 2 Derived Reactive Oxygen Species Promote CD8^+^ T cell Effector Function": 21-00180-FL-Supplement

**Supplemental Table. Sequences of primers used for detection of target cDNA.**

| Target | Forward Primer Sequence | Reverse Primer Sequence |
|---|---|---|
| CD107a | 5′-CTATCCAGGCCTACCTGTCG | 5′-GGATACAGTGGGGTTTGTGG |
| FasL | 5′-ACTCCGTGAGTTCACCAACC | 5′-TTAAATGGGCCACACTCCTC |
| Granzyme B | 5′-CTGACCTTGTCTCTGGCCTC | 5′-CTCTCGAATAAGGAAGCCCC |
| IL-2 | 5′-TCCTGAGCAGGATGGAGAAT | 5′-GTCAAATCCAGAACATGCCG |
| IL-4 | 5′-TCTCGAATGTACCAGGAGCC | 5′-GGTGTTCTTCGTTGCTGTGA |
| IFNγ | 5′-AGGCCATCAGCAACAACATA | 5′-TGAGCTCATTGAATGCTTGG |
| Perforin | 5′-AAGGTAGCCAATTTTGCAGC | 5′-CTGAGCGCCTTTTTGAAGTC |
| TGFβ | 5′-GGAGAGCCCTGGATACCAAC | 5′-AAGTTGGCATGGTAGCCCTT |
| TNFα | 5′-GGTCTGGGCCATAGAACTGA | 5′-CAGCCTCTTCTCATTCCTGC |
| GAPDH | 5′-TTGATGGCAACAATCTCCAC | 5′-CGTCCCGTAGACAAAATGGT |
| β-globin | 5′-GAAGCGATTCTAGGGAGCAG | 5′-GGAGCAGCGATTCTGAGTAGA |

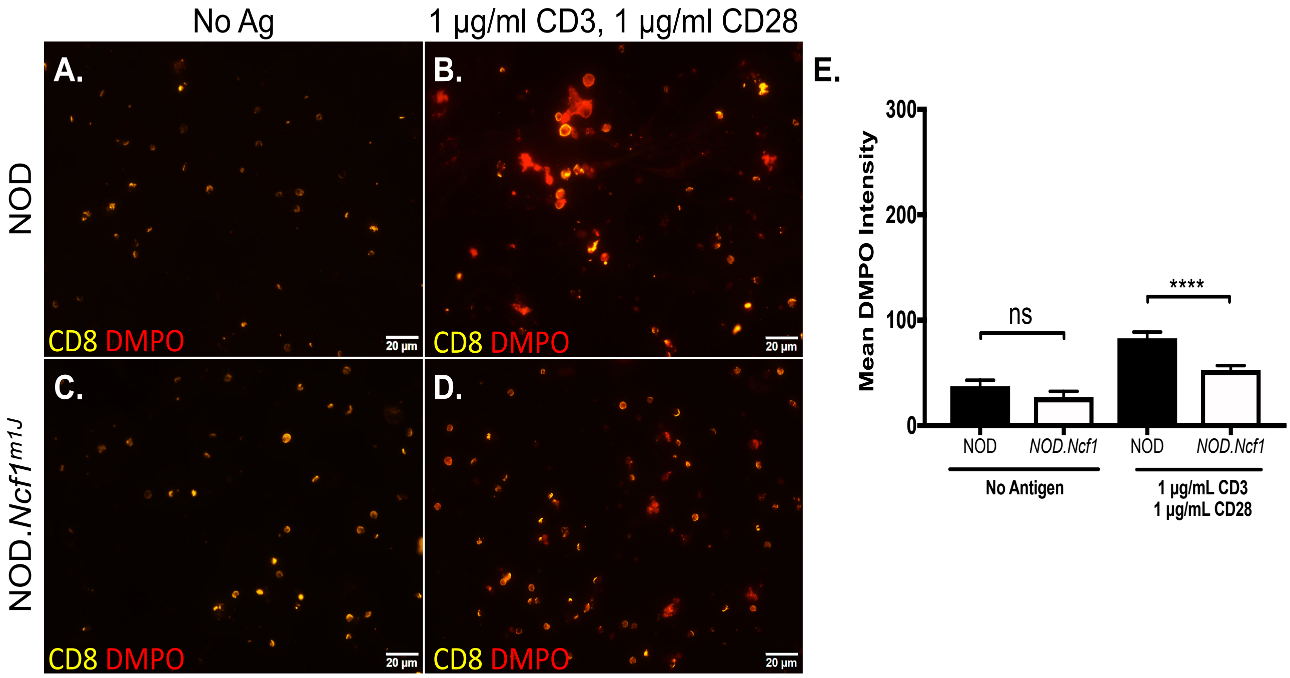


**Supplemental Figure 1. NOD.*Ncf1$^{m1J}$* CD8$^+$ T cells exhibit restrained ROS production after polyclonal stimulation.** CD8$^+$ T cells from NOD or NOD.*Ncf1$^{m1J}$* mice were stimulated with α-CD3 and α-CD28 in the presence of the spin trap DMPO. DMPO adducts were observed from (A) unstimulated NOD CD8$^+$ T cells, (B) stimulated NOD CD8$^+$ T cells, (C) unstimulated NOD.*Ncf1$^{m1J}$* CD8$^+$ T cells, or (D) stimulated NOD.*Ncf1$^{m1J}$* CD8$^+$ T cells using an Olympus IX81 Inverted Microscope with a 60x objective. Images were analyzed using ImageJ and give a pseudo-color: yellow for αCD8 and red for DMPO. Statistical comparisons were made amongst the means of the unactivated or stimulated T cells after 24 hours (**** $P \leq 0.0001$).

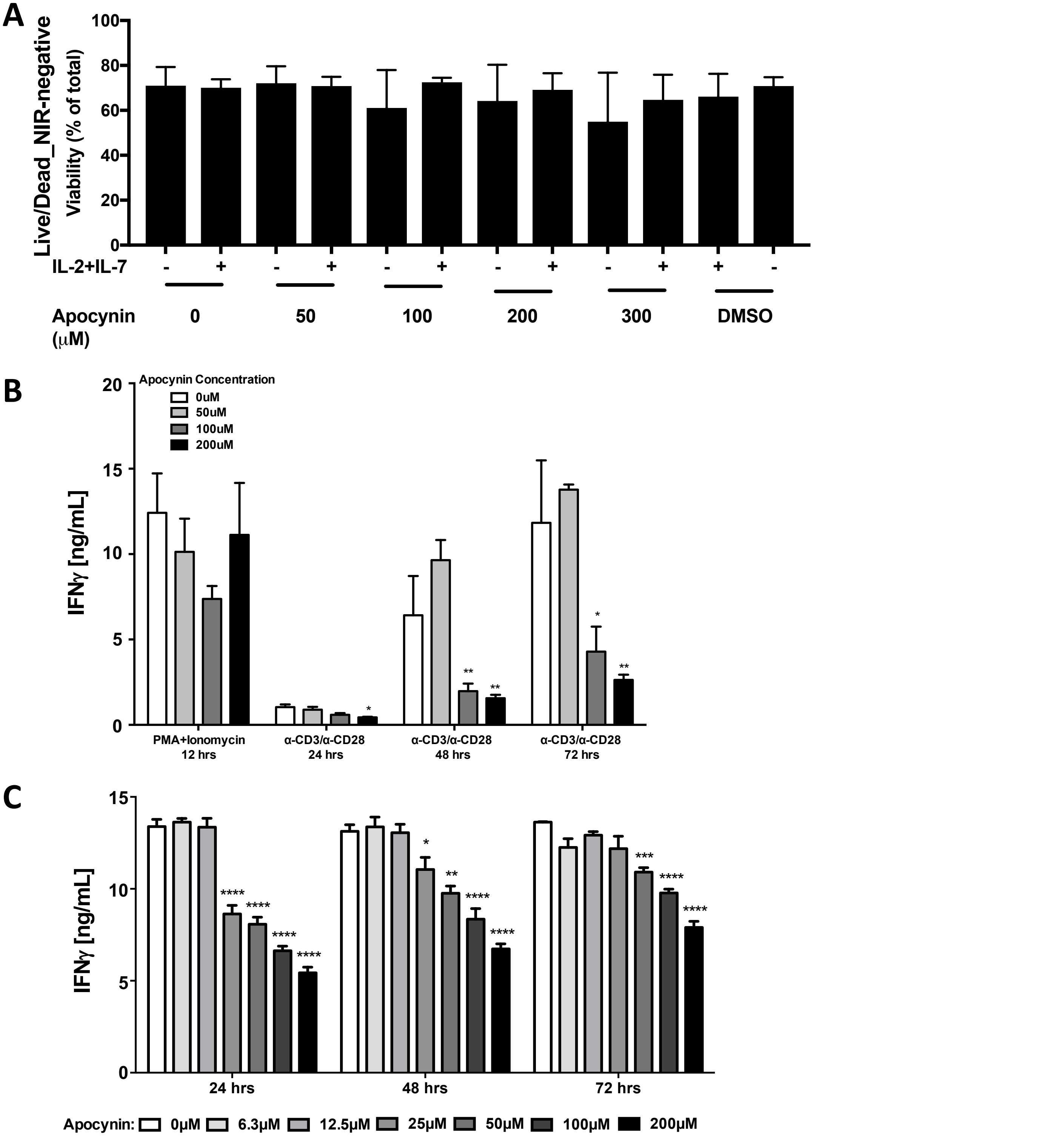


**Supplemental Figure 2. Dose Dependent Repression of IFNg production by Apocycin in mouse and human CTL.** CD8+ T cells from NOD mice were stimulated with α-CD3 and α-CD28 conjugated beads in the presence of the indicated concentrations of apocynin. (A) Viability was measured by flow cytometry using live-dead near infrared staining at 72 hrs. (B) IFNγ was measured by ELISA in supernatants at the indicated times. (C) CD8+ T cells from healthy volunteers were stimulated with α-CD3 and α-CD28 conjugated beads in the presence of the indicated concentrations of apocynin. IFNγ was measured by ELISA in supernatants at the indicated times. Statistical comparisons were made for the means of each dose versus the untreated group for the same time point. (* $P \leq 0.05$; ** $P \leq 0.01$; *** $P \leq 0.001$; **** $P \leq 0.0001$).

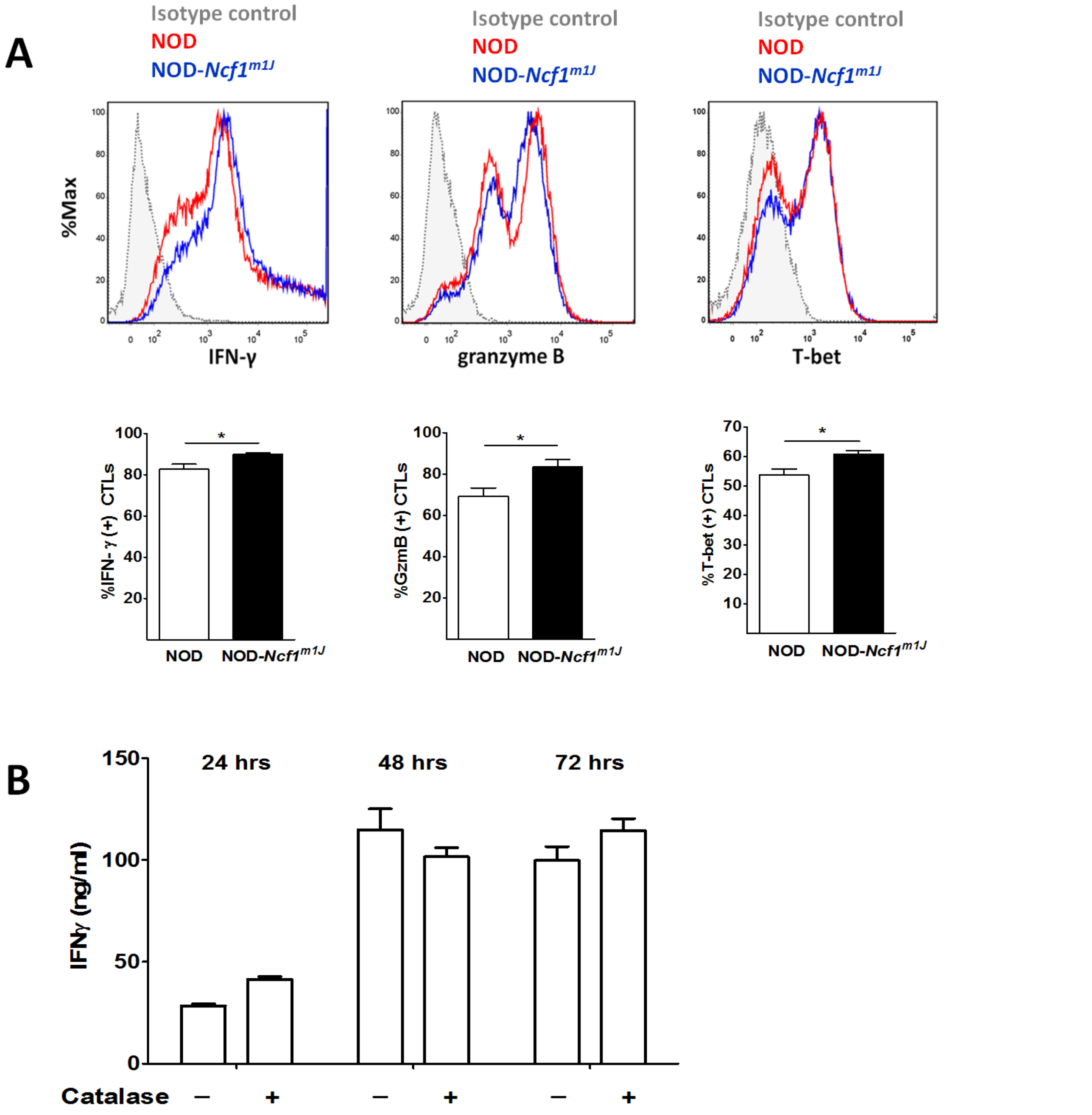


**Supplemental Figure 3. NOX2-derived ROS are not required for effector molecule production by CD8$^+$ T cells when activated by PMA and ionomycin.** (A) Histogram and the quantitation of intracellular staining in purified CD8$^+$ T cells (5 × 10$^5$ cells) after 50ng/mL PMA and 1μg/mL ionomycin stimulation for 48 hours with PMA/ionomycin plus Golgi stop added for the final 6 hours. At least four mice were included in each group. (B) NOD CD8$^+$ T cells were stimulated by PMA/ionomycin in the presence or absence of 1000U/mL catalase. After 24, 48, and 72hrs, supernatants were collected, and IFNγ was measured by ELISA. Results are presented as the means (±SEM). Results are compared using Student's t test (* $P \leq 0.05$).
